# Umbilical cord structure shapes feto-maternal heat transfer across mammals

**DOI:** 10.1101/2025.11.18.688874

**Authors:** Tianran Wan, Davis Laundon, Shier Nee Saw, Nicholas Cheng, Hwee Kuan Lee, Edward D. Johnstone, Oliver E. Jensen, Rohan M. Lewis, Igor L. Chernyavsky

## Abstract

The umbilical cord varies widely across placental mammals, yet the functional consequences of this structural diversity remain unclear. We compared cord anatomy across 130 mammal species, reconstructed ancestral states, and combined multimodal imaging with physics-based modelling of oxygen and heat transfer. Reconstructions indicate that the ancestral umbilical cord was uncoiled, with three vessels and an allantoic duct; the human combination of coiling with loss of the duct is rare. Across species, vessel number, duct persistence, and cord length increase with birth weight. Modelling predicts that oxygen delivery remains near maximal across architectures, but that placental heat loss depends more strongly on umbilical flow and on cord length and coiling, suggesting more efficient cooling in larger mammals. Microscopy and ultrasound imaging reveal geometric vessel layouts, including a human triple helix, consistent with configurations predicted to reduce inter-vessel heat shunting. As fetuses must dissipate heat through the mother, these results link cord morphology, evolution, and fetal thermal resilience.

## Introduction

The mammalian placenta is a transient organ responsible for physiological exchange of respiratory gases, nutrients and metabolites between mother and fetus during gestation. Although placental mammals share conserved placental functions and a common evolutionary origin, they display a striking diversity of placental structures across physiological scales (*1, 2*). The evolutionary drivers of this diversity remain unresolved, with proposed factors including reproductive strategy (*3*), environmental and parasite pressures (*4*), and feto-maternal conflict (*5, 6*). What is less appreciated is that the umbilical cord displays an equally remarkable level of structural variation across placental mammals. Computational modelling has recently clarified how placental architecture shapes transport (*7, 8*), yet the structure–function relationship of the mammalian umbilical cord, particularly across species and in atypical human pregnancies, remains largely unexplored.

The umbilical cord is the fetus’ vital conduit to the placenta at the maternal vascular interface, delivering oxygen and nutrients via the umbilical vein and removing waste via the umbilical arteries (Fig. 1). The cord structure varies widely among mammals in vessel number, the presence of an allantoic duct, and the degree of spiralling (*1, 9*). Despite this diversity, the cord’s comparative anatomy, and the functional implications of this variability, have received even less attention than the placenta.

**Figure 1.**
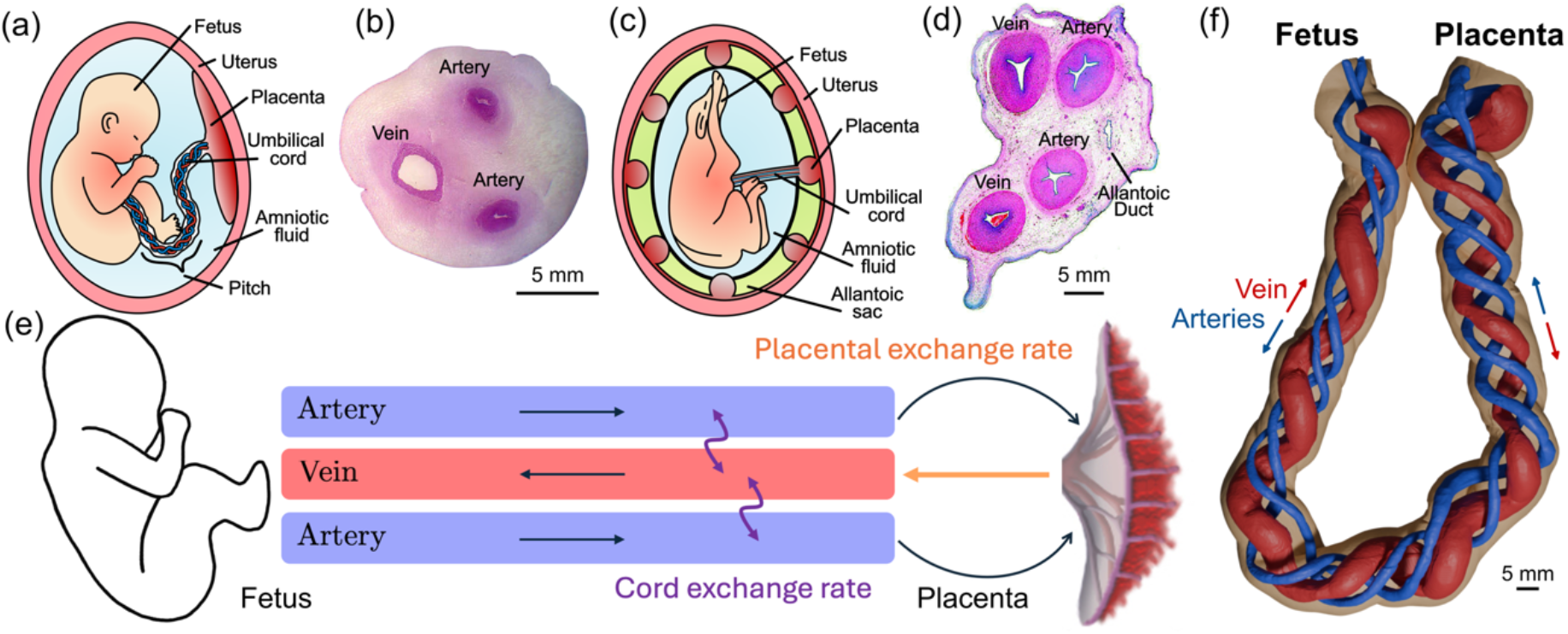
Architecture of the mammalian umbilical cord: **(a)** Schematic of the helical, 3-vessel human umbilical cord connecting the fetus to a discoid placenta; the pitch represents the average length of a coil normalized by the cord’s radius (reciprocal of the cord helicity, 2π/*Ω*, and umbilical coiling index, 1/UCI). **(b)** Histological cross-section of a healthy human cord (adapted from (*56*) under CC BY-NC 4.0). **(c)** Schematic of a cotyledonary cow placenta with an unspiralled, 4-vessel cord. **(d)** Histological cross-section of the cow umbilical cord, showing an allantoic duct (absent in typical human cords; adapted from (*45*), with permission). **(e)** Diagram of counter-current exchange in a 3-vessel cord, indicating flow directions for each vessel between fetal and placental ends. **(f)** Three-dimensional micro-CT reconstruction of umbilical vessels in a normal human cord, highlighting their triple-helical geometric arrangement (with no single vessel serving as a central ‘axis’).

Thermal regulation is a critical and underappreciated cord function. Fetal temperature typically exceeds maternal temperature by about 0.5 – 1 °C (*10*) due to high metabolic heat production (mass-specific heat production in the fetal lamb is approximately double that of the adult (*11*), ranging from about 1 W for rodents to 1 kW for large mammals (*12*)). Unlike adults, which regulate temperature via the skin and evaporative cooling, the fetus relies primarily on placental and umbilical blood flow for heat exchange (*13, 14*). Although prior studies have highlighted the importance of cord structure in feto-maternal heat transfer (*15, 16*), the consequences of inter- and intra-species structural diversity for fetal thermal regulation remain largely unknown.

Across mammals, umbilical cords can contain two vessels (one vein, one artery), three vessels (one vein, two arteries), or four vessels (two veins, two arteries). The allantoic duct accompanies the vessels in most species, connecting the fetus to the allantoic sac (Fig. 1); it is reduced or absent in rodents and becomes obliterated in primates by mid-gestation (*17*). Similarly, human embryonic cords start with four vessels, with the right umbilical vein regressing by about the 6^th^ week of gestation (*18*). Spiralling is a hallmark of the human umbilical cord (Fig. 1f) but is uncommon in most other mammals. How these features shape solute transport and heat exchange is unclear. In particular, diffusion between closely-spaced umbilical arteries and vein(s), carrying counter-current flows, creates the potential for a functional shunt in feto-placental exchange (*15*): for oxygen, oxygenated venous blood could lose some oxygen back to the arteries before reaching the fetus; for heat, excess arterial fetal heat could return via the umbilical vein(s). The extent of such diffusive coupling depends on the cord’s vascular anatomy and haemodynamics.

Parallel variation occurs within humans. Most human cords have two arteries and one vein, but a single umbilical artery (SUA) occurs in a minority of pregnancies (*19*), and rare four-vessel cords have been described (*20*). Remnants of the allantoic duct may persist in as many as 15% of cases (*21*). The degree of coiling can be quantified by the umbilical coiling index (UCI), defined as the number of coils per unit length of cord (*22*), with hypo- and hyper-coiled cords representing 13% and 21% of all singleton pregnancies respectively (*23*). How deviations from the typical three-vessel moderately-coiled human cord influence its function is poorly understood.

The structural diversity of the umbilical cord across mammals and within human pregnancies offers a unique opportunity to unravel the interplay between structure and physiological function. Analysis of the structure of the umbilical cord in different species provides insight on how such variation influences the evolution of mammalian reproductive strategies as well as shedding light on the foundations of healthy human pregnancies and the pathological consequences of atypical cord structure.

Here we integrate comparative anatomy and evolutionary perspectives with mathematical modelling to examine how umbilical cord structure influences fetal oxygen supply and thermal regulation. We analyse umbilical vessel number, arrangement and coiling across a wide range of placental mammals and in atypical human cords, and we develop a model of solute and heat transfer to quantify how the cord’s structural variation may affect transfer efficiency across species and pregnancy characteristics. Our theoretical framework is informed and validated by multi-modal *in vivo* and *ex vivo* imaging.

## Results

### Reconstructing the origins of structural diversity of the umbilical cord in mammals

Ancestral trait reconstruction across *n* = 130 placental mammal species (Fig. 2) revealed that the ancestral umbilical cord had three vessels (root node probability: two = 0%, three = 97.5%, four = 2.5%), an allantoic duct (97.6%), and was likely unspiralled (82.4%). The human umbilical cord structure is almost unique among mammal configurations, having lost the allantoic duct and acquired a spiralled structure as later innovations, while retaining the ancestral three vessels. This configuration represents only 3% of the species in our dataset (all great apes or Old World monkeys).

**Figure 2.**
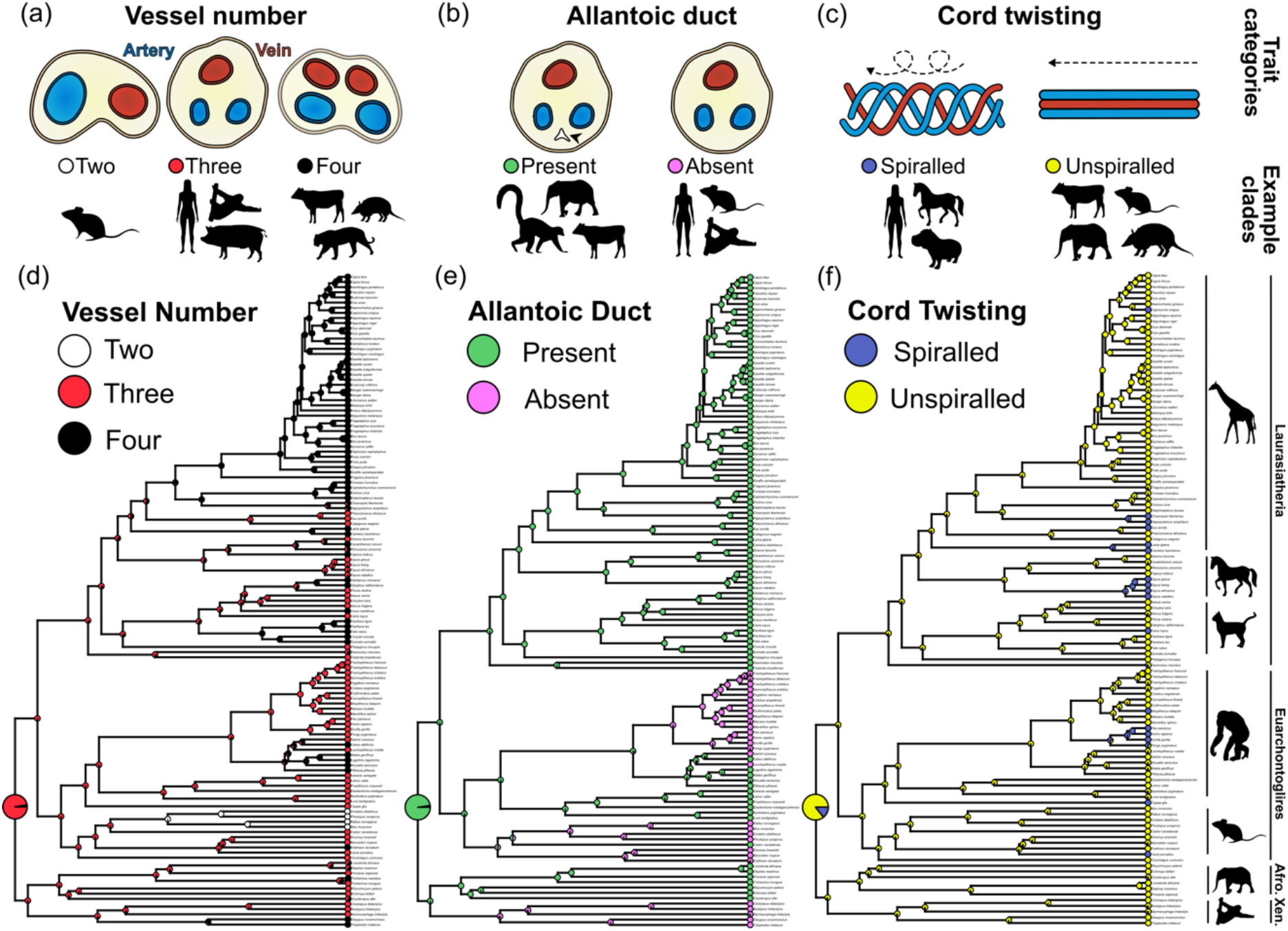
Evolution of umbilical cord architecture: **(a-c)** Diagrammatic summaries of structural trait categories (top) with example placental mammal clades where these traits are widespread (bottom): vessel number (a), presence of the allantoic duct (b), and vascular spiralling status (c). **(d-f)** Ancestral-state reconstructions for umbilical cord traits across the placental mammals. The inferred ancestral cord had three vessels (d), an allantoic duct (e), and was likely unspiralled (f). (Animal silhouettes dedicated to the public domain were adapted from www.phylopic.org.)

Three-vessel cords are predominantly found in catarrhine primates (Old World monkeys and great apes), strepsirrhine primates and kin (lemurs, tarsiers, colugos), non-myomorph rodents (beavers and guinea pigs), lagomorphs (rabbits), tenrecs, pilosan xenarthrans (anteaters and sloths), suids (pigs), equids (horses, with occasional 3–4-vessel branching), and some carnivorans. We infer a single transition to a 2-vessel cord in muroid rodents (mice, rats, hamsters). Independent transitions to a 4-vessel cord occurred at least five times: in artiodactyls (‘even-toed ungulates’, including cows, antelopes, dolphins), pinnipeds (walruses and sea lions), feliform carnivores (cats, hyenas, meerkats), platyrrhines (New World monkeys), and cingulates (armadillos). Whether pigs reverted from a 4-vessel artiodactyl ancestor to a 3-vessel cord, or camelids (camels and llamas) independently evolved to four vessels within artiodactyls, remains unresolved.

The allantoic duct was lost at least three times independently: in catarrhine primates; rodents (with uncertainty as to whether muroids lost the duct independently); and in xenarthrans (sloths, anteaters, armadillos). An unspiralled cord is the norm across placental mammals. However, spiralling evolved at least four times independently, in hippos, camelids, horses, and great apes. Transition counts are conservative because we excluded changes represented by single terminal taxa; denser sampling will be needed to fully resolve those cases. Together, these reconstructions underscore the evolutionary distinctiveness of the human cord and motivate the functional analysis presented below.

### Structure and heat transfer efficiency of the cord correlate with birth weight

Across species, umbilical cord length scales with birth weight with an exponent near 1/3 (R^2^ = 0.49, *p* < 0.001; Fig. 3a). Relative to this trend, uncoiled cords are shorter and coiled cords are longer for a given birth weight; the cord length–birthweight relationships differ between coiled and uncoiled cords (ANCOVA, *p* < 0.05; Fig. 3a). Species with 4-vessel cords give birth to larger young than species with 3-vessel cords (*p* < 0.001), whereas 2-vessel cords were observed only in muroid rodents with very small neonates (Fig. 3b). Species with an allantoic duct have higher birth weights than those without (*p* < 0.001). We did not detect a birthweight difference between spiralled and unspiralled cords. Similar patterns hold when using litter size, gestation time, litter investment (litter size × birth weight), or a precociality metric ([birth weight × gestation time] / litter size) (Supplement, Fig. S3).

**Figure 3.**
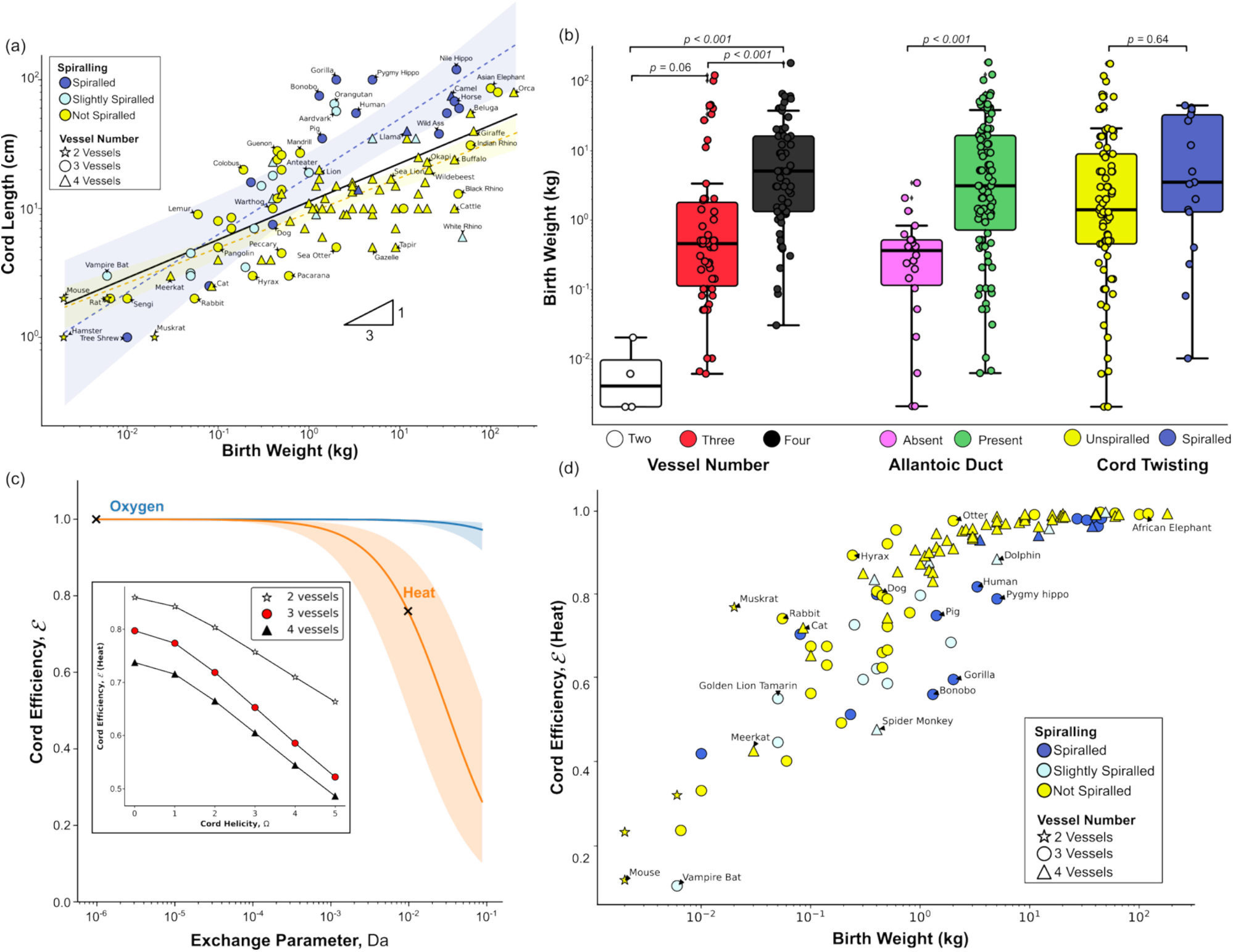
Allometric scaling and oxygen/heat transfer efficiency of the umbilical cord: **(a)** Scaling of cord length (cm) with birth weight (kg) across species. Colours represent degree of cord spiralling; point shapes indicate vessel number; lines show linear regressions for the whole dataset (solid black; R^2^ = 0.49), coiled cords (dashed blue; R^2^ = 0.69), and uncoiled cords (dashed orange; R^2^ = 0.58); shaded areas are 95% confidence intervals. **(b)** Comparison of cord trait categories (vessel number, allantoic duct presence, spiralling status) by birth weight (kg). **(c)** Predicted cord efficiency ℰ(Methods, Eq. 1) as a function of Da for a symmetric 3-vessel human cord; for oxygen, the exchange parameter Da ∼ 10^−6^ and the normalized placental flux *N*_*p*_ ≈ 0.03 (range: 0.01 – 0.1; shaded blue); for heat, Da ∼ 10^−2^ and *N*_*p*_ ≈ 3 (1 – 10; shaded orange). Increased helicity raises the normalized inter-vessel exchange flux *N*, which lowers ℰ for heat. Inset: Predicted ℰ for heat versus helicity for cords with different number of vessels. **(d)** Predicted ℰfor heat versus birth weight (kg) across species, using allometric scaling for cord flow rate and length. Larger species approach ℰ ≈ 1, indicating more efficient placental dissipation of fetal heat.

We quantify the cord transfer efficiency ℰ (defined in Methods, Eq. (1), and Supplement) for oxygen and heat across 2-, 3- and 4-vessel configurations, while varying helicity, cord length and flow. For oxygen, the characteristic exchange parameter Da (a dimensionless Damköhler number) is very small (≈10^−6^; see Supplement) and ℰ approaches 1 for all cord architectures (Fig. 3c), ensuring near-perfect oxygen delivery to the fetus across species. For heat, however, Da for the human cord is typically 10^−2^, giving ℰ ≳ 0.5. For abnormally long cords or with reduced umbilical flow, Da can approach 10^−1^; higher helicity (hypercoiling) further increases inter-vessel exchange *N*, lowering ℰ.

In that regime, only up to about half of the additional fetal heat is removed by the human placenta. ℰ also declines modestly as the number of umbilical vessels increases at fixed cord length *L* and umbilical flow rate *Q* (Fig. 3c, inset). The helicity range tested corresponds to typical human umbilical coiling indices (UCI ≈ 0.1 coils/cm; see Methods) and overlaps equine measurements.

Allometric scaling suggests that larger species have more efficient cord-mediated heat removal. From the limited literature with umbilical flow measurements (in human, sheep, pony, guinea pig) (*24*–*27*), we fit umbilical flow rate (cm^3^/s) vs. birth weight (kg), using *Q* ≈ 1.7 BW^α^, with α ≈ 1.0 ± 0.14, consistent with major artery flow scaling (*28, 29*). Using this scaling we estimate *Q* across placental mammals. Our model predicts that ℰ for heat increases non-linearly with birth weight (Fig. 3d; Methods, Eq. 1). Larger species approach higher efficiencies (up to ℰ ≈ 1 for cetaceans and elephants), indicating more efficient dissipation of fetal heat via the placenta, while smaller or litter-bearing species have significantly lower ℰ.

### Cross-sectional cord configurations typically minimize inter-vessel exchange

We mapped the normalized cross-sectional heat exchange flux *N* between the umbilical vessels as a function of vascular configuration parametrized by the angle between a vein and an artery *θ*_**1**_ (measured relative to the line passing through the centres of the cord and the vein) and the inter-arterial angle Δ*θ* (Fig. 4a-c). In simulations, vein positions were fixed while relative arterial positions were varied.

**Figure 4.**
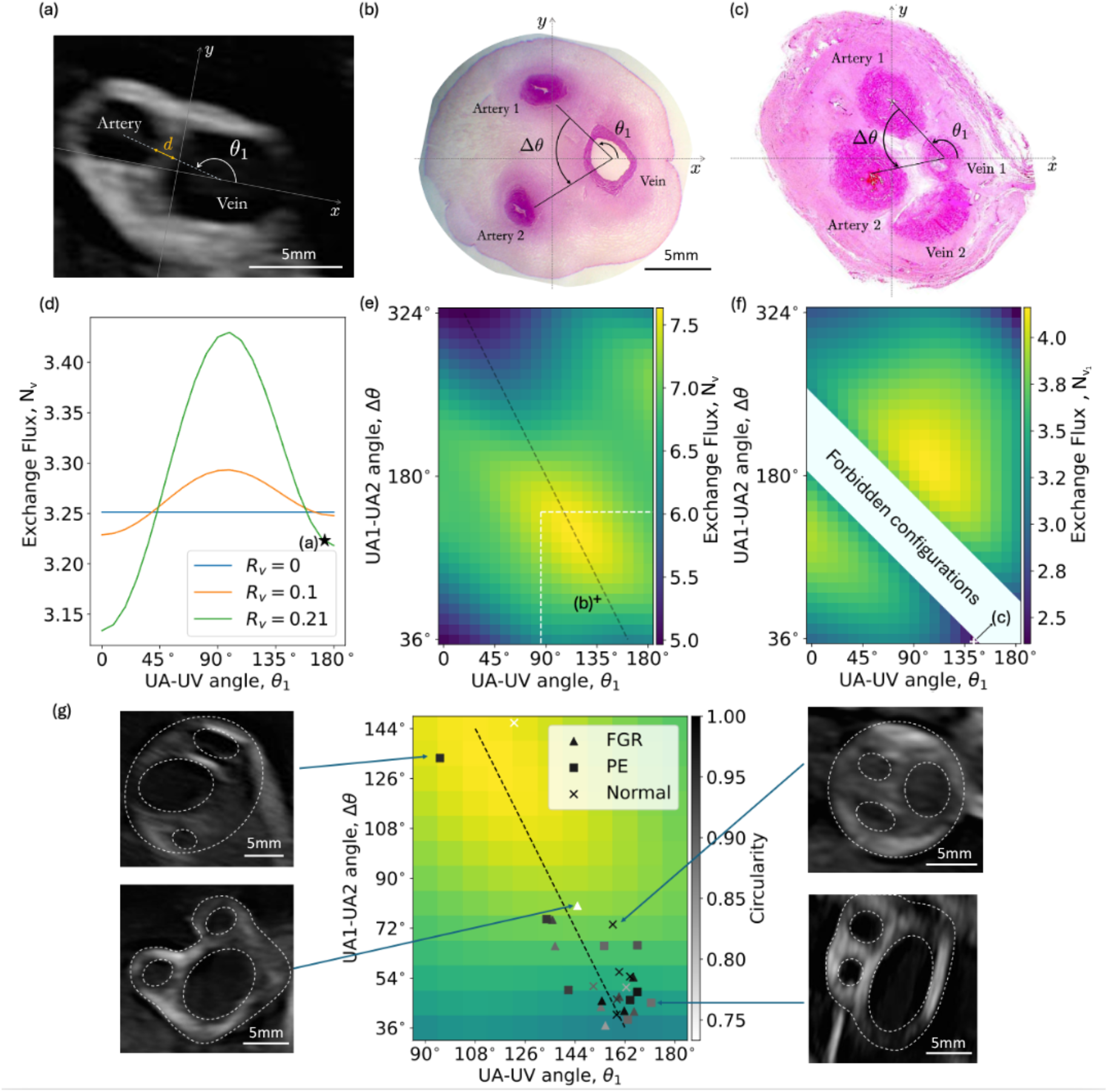
Cross-sectional cord configurations and inter-vascular exchange: **(a)** Ultrasound cross-section of a cord with a single umbilical artery (SUA) from a pre-eclamptic (PE) pregnancy; *θ*_**1**_ is the angle between the vein and the artery relative to the *x*-axis, connecting the centre of the cord to the centre of the vein; *d* is the artery–vein separation distance normalized by the cord radius. **(b)** Histological cross-section of a healthy human 3-vessel cord (adapted from (*56*) under CC BY-NC 4.0); the cord configuration is parametrized by *θ*_**1**_ and the inter-arterial angle Δ*θ*. **(c)** Histological cross-section of a hippopotamus 4-vessel cord (adapted from (*45*), with permission); the geometry is parametrized by *θ*_**1**_, Δ*θ* and the inter-venous angle *θ*_v_. **(d-f)** Predicted configuration maps for the normalized cross-sectional exchange flux *N*_v_ between one vein and the artery(ies): 2-vessel cords with different vein offset *R*_v_ (d; star marks the configuration in (a)); 3-vessel cords (e; black cross marks the configuration in (b); dashed line is the symmetry line Δ*θ* = −2*θ*_**1**_ + 2π), and 4-vessel cords (f; white cross marks the configuration in (c)). Lower *N*_v_ indicates reduced diffusive coupling between vessels; for 2-, 3-vessel cords, the total exchange flux *N* = *N*_v_ (see Supplementary Tables S1-S3 for more details). **(g)** Exchange flux map (delineated by the white dashed lines in (e)) with angle measurements from *in vivo* ultrasound in normal, fetal growth restricted (FGR), and PE pregnancies; most cords cluster near low-*N*_v_ regions. Representative ultrasound cross-sections are shown.

For a 2-vessel cord, *N* is maximized when the artery is transversely orthogonal to the cord–vein axis (*θ*_**1**_ ≈ *π*/2). Notably, a representative single umbilical artery (SUA) configuration of atypical human cord *in vivo* (Fig. 4d; *θ*_**1**_ ≈ *π*) lies near a local minimum of *N*. For a 3-vessel cord, the extrema of *N* lie on the symmetry line Δ*θ* = –2*θ*_**1**_ + 2*π* (Fig. 4e). Along this line, flux out of each artery is half the venous flux (see Supplement). Finally, for a 4-vessel cord, the minimum-*N* configuration also corresponds closely to observed histological cross-sections *ex vivo* (Fig. 4f; see Supplement for more details).

*In vivo* ultrasound imaging of umbilical cross-sections from normal and pathological human pregnancies showed that most cords cluster near the symmetry line Δ*θ* = –2*θ*_**1**_ + 2*π* and in low-*N* regions (Fig. 4g). There was no significant difference between groups. Rare high-*N* configurations, with arteries flanking the vein transversely, occurred in both groups. Cross-sectional morphometrics (Supplement, Fig. S4) showed a reduced Wharton’s jelly fraction and a relatively enlarged venous area in pre-eclamptic (PE) pregnancies (p < 0.001; see Methods), with no groupwise differences in scaled vein offset or artery–vein separation. Given the small sample size, we cannot infer clinical significance.

## Discussion

The mammalian umbilical cord shows striking structural diversity; however, until now the physiological consequences of this variation were not well understood. By integrating comparative anatomy, phylogenetic reconstruction and physiological modelling with *in vivo* and *ex vivo* imaging, we identify three broad conclusions. First, the ancestral placental mammal likely had an uncoiled, 3-vessel cord with an allantoic duct. The human umbilical cord configuration is almost unique among mammals in retaining three vessels while acquiring coiling and losing the allantoic duct. Second, cord architecture is aligned with birth weight (consistent with allometric scaling (*29, 30*)): vessel number, allantoic duct presence and cord length all correlate with neonatal mass, and our mathematical model predicts that oxygen delivery remains near-maximal across cord architectures whereas heat-removal efficiency is more sensitive to blood flow, cord length and coiling. Third, cords across placental mammals tend to adopt cross-sectional vascular configurations that minimize inter-vessel exchange, consistent with reduced vascular shunting that could undermine fetal thermal regulation.

Competing hypotheses for the origin of coiling propose that it arises from fetal movement or from intrinsic cord properties (e.g., asymmetric blood flow or microstructural anisotropy) (*9, 31*–*33*). Our comparative analysis shows that coiling is rare across mammals yet prominent in humans and some ungulates. Between species, longer cords tend to be more coiled, which is consistent with the fetal-movement hypothesis. Equine observations that cord twisting can be mechanically unwound (*34*), correlation of restricted fetal activity with shorter cords (*35, 36*), and association of hypo-/hyper-coiled cords with adverse clinical outcomes (*22, 37*) further support the interpretation of coiling as a function of fetal movements and cord length. Our theoretical modelling predicts that coiling modestly reduces feto-placental heat-removal efficiency at typical human coiling indices; however, efficiency can decline more substantially in long or hypercoiled cords, especially when umbilical flow is reduced, which is also indicated in animal studies (*38, 39*).

Across species, umbilical flow rate scales approximately linearly with birth weight, while the metabolic heat-generation rate scales sublinearly (*12, 30*). As a result, our mathematical modelling predicts that heat-removal efficiency increases with neonatal mass, approaching unity in the largest species. This pattern aligns with life-history strategies: large-bodied mammals with singleton precocial offspring (e.g., many ungulates, cetaceans, elephants) tend to achieve higher predicted heat-removal efficiency, whereas species with altricial young or larger litters (e.g., dogs, rabbits, mice) show lower efficiency. Comparative anatomy echoes this pattern: 4-vessel cords and allantoic ducts are more common in species with heavier neonates, consistent with greater maternal investment in umbilical flow capacity, thermal buffering and fetal resilience.

Our simulations predict that certain umbilical vascular configurations minimize inter-vessel heat exchange, which matches observations in *ex viv*o histology and *in vivo* ultrasound imaging from both normal and atypical human cords (consistent with our prior study in healthy cords (*15*)). Transversely symmetric configurations that minimize shunting are typical for a triple helix, thus challenging the paradigm in which umbilical arteries are twisted around a straight umbilical vein. Notably, representative cords with a single umbilical artery and mammalian cords with four vessels also lie near local minima of predicted shunting. Our findings indicate that the vascular architecture of mammalian cords supports feto-placental heat transfer and limits counter-current cross-vessel shunting.

Overall, fetal thermoregulation depends on utero-placental and umbilical blood flows. Maternal temperature variations are mirrored by the fetus, whereas fetal temperature changes do not measurably affect the mother (*14*). Our results imply that long cords, hypercoiling or reduced umbilical flow could compromise fetal cooling capacity, heightening the risk of fetal hyperthermia during maternal fever or extreme heat exposure, with both conditions being associated with adverse pregnancy outcomes and developmental disorders in humans (*40, 41*). Because the cords of smaller mammals and, by extension, of smaller-bodied human fetuses have lower heat-removal efficiency, increased environmental heat may pose disproportionate risks. Conflicting reports of the impact of heat stress on utero-placental flow in animals (*25, 42*) and birth weight in human pregnancies (*41*) underscore the need for direct, species-specific measurements of feto-placental flow and heat transfer under thermal challenge in future studies.

Our theoretical framework captures dominant structure–function relationships in the mammalian cord; however, it does not resolve species-specific placental microanatomy or active vascular regulation. Flow data across mammals are sparse, and phylogenetic cord anatomy was not available for some taxa, so feto-placental heat-transfer efficiency predictions should be viewed as qualitative trends rather than precise estimates. Future work should pair species-specific cord and placental architectures with umbilical and uterine flow measurements, explore variations in cord geometry across gestation and test predicted vulnerabilities of feto-maternal heat transfer experimentally *in vivo* and *ex vivo*, e.g., using population-wide studies or biomimetic approaches.

Beyond thermal regulation, multiple other functional constraints exist. For example, human cords *ex vivo* show complex deformation–flow coupling augmented by Wharton’s-jelly poroelasticity (*43, 44*). Although the observed cord’s structure is consistent with reduced thermal shunting, the role of cord structure in the mechanical stability of the umbilical circulation remains an open question.

Our study reframes the umbilical cord as a heat-transfer system. It proposes a new paradigm in which the triple helix of the human cord, and the vascular architecture of the cords in other placental mammals, may help support fetal thermal regulation by minimizing shunting. The identified structure–function relationships help explain evolutionary patterns that encompass both normal and atypical human cord phenotypes, pointing to testable hypotheses for pregnancy monitoring and for understanding how environmental heat stress may differentially impact fetal development across mammals.

## Materials and Methods

### Data Collection

Data for umbilical cord characteristics and life history strategy metrics (i.e. birth weight, gestation time, litter size) were compiled from Benirschke’s online Comparative Placentation resource from the University of California San Diego (*45*). Only species with complete information on vessel number, duct presence, cord spiralling, and term length were included, resulting in a final dataset of 130 placental mammal species. The full dataset is available (see Supplement for repository). Where additional sources have been used for a species beyond the online resource (*45*), these are also listed.

Ultrasound images of cord cross-sections were obtained at St Mary’s Hospital, Manchester, UK, with written informed consent and ethical approval from the North West – Greater Manchester Research Ethics Committee (15/NW/0829). Ultrasound measurements of the cord were performed on GE Voluson E10 (GE Healthcare, Chicago, IL, USA) during routine clinical visits (gestational age between 20-34 weeks). The control group included 9 individuals with normal estimated fetal weight (EFW). The pathological group consisted of 10 participants with fetal growth restriction (FGR), defined as EFW below the 3^rd^ centile for gestation, and of 10 participants with pre-eclampsia (PE).

A semi-automated Python script (see Supplement) was used to extract structural metrics from the images, including artery-vein separation distances, vein offset from the cord’s centre, characteristic angles, the area of each umbilical vessel and Wharton’s jelly area (cord area excluding vessels area). All measurements were normalized by the cord’s radius.

### 3D imaging of human umbilical vessels

To generate the 3D image of human umbilical vessels shown in Fig. 1f, an entire placenta and its cord were obtained from an uncomplicated delivery from a normal term pregnancy with written informed consent and ethical approval from the Southampton and Southwest Hampshire Local Ethics Committee (11/SC/0529). The placenta and cord were fixed in 10% formalin for a week, stained in 10% Lugol’s iodine solution, mounted to a polystyrene board, and imaged in a diondo d5 microCT system as described previously (*46*). Vessels were segmented in Microscopy Image Browser (*47*) and visualized in Blender v.4.1.

### Ancestral trait reconstruction

Ancestral umbilical cord characteristics were reconstructed using stochastic character mapping and likelihood modelling using the R packages *phytools* v.2.1.1 (*48*) and *APE* v.5.8 (*49*), run with R v.4.4.0 implemented in RStudio v.2024.4.1.748. Phylogenetic trees were built for the mammal species compiled above based on the phylogeny outlined by Upham *et al*. (*50*) using the online resource (*51*). Consensus trees were compiled from 1,000 tree replicates using the *consensus*.*edges* function in *phytools*. Occasional missing data values for certain traits in certain species (see Supplement) were excluded from ancestral trait reconstruction in their respective analyses, so the topology of each tree is very slightly different. Practically speaking, these phylogenies can be considered as one. Traits were coded as a matrix and, where there was ambiguity in a species’ trait categorisation for ‘intermediary’ descriptions (i.e. 3 *or* 4 vessels, a ‘remnant’ allantoic duct, or a ‘slightly’ spiralled cord), then these traits were coded as having an equal probability of representing either end of the trait binary. For each trait, the fit of different variations of the extended M*k* model (*52*) (ER: equal rates, SYM: symmetric rates, and ARD: all-rates-different, with both Fitzjohn and MCMC root priors) to the data was compared by calculating Akaike Information Criterion (AIC) scores and weights for each model. We simulated 1,000 stochastic character maps for the evolution of each trait, weighting the use of each model variation into the simulations as calculated above. Trait changes and internal node likelihood were then calculated as a summary from across the 1,000 maps.

### Traits by life history metrics

Linear regression of cord length on birth weight was fitted using log-log transformed data. For each species, cord variation was compared against birth weight, litter size, and gestation time. For each trait, data was assessed for normality and homogeneity of variance, with Shapiro and Levene’s tests respectively, and an appropriate non-parametric or parametric test selected. Cord vessel numbers were compared by birth weight using a Kruskal-Wallis test followed by a Dunn’s *post hoc*, and allantoic duct traits and spiralling were assessed using a Mann-Whitney-U test and *t*-test respectively. All statistical comparisons were conducted using the Python package *SciPy* v. 1.11.4 (*53*) run with Python 3.11.7 implemented in Jupyter Notebook 7.0.8.

### Counter-current solute and heat transfer model

The umbilical cord is modelled as a straight circular cylinder containing *n* arteries and *m* veins within its cross-section in the plane orthogonal to the axis of the cylinder, extending the theoretical framework for three helical vessels presented in (*15*). Each vessel has a constant and common pitch (the length of a complete helical turn) of *2π/Ω^∗^*, where *Ω*^∗^ is the parameter describing helicity, proportional to the umbilical coiling index, UCI = *Ω^∗^/2π* coils/cm (see Supplement for more details). In normal pregnancies, the mean UCI is 0.17 coils/cm (*22*); cords with UCI below the 10th centile (0.07 coils/cm) are classified as hypocoiled while cords with UCI above the 90th centile (0.3 coils/cm) are classified as hypercoiled.

The exchange of solute or heat between the cord’s vessels is modelled as a steady diffusion process in the cord tissue, coupled with cross-sectionally averaged transport along the umbilical arteries and veins. For simplicity, we assume no flux across the outer surface of the cord (which is shown to have a modest effect (*15*)) and an azimuthally uniform but axially varying concentration, or temperature, *c*^*^ along each umbilical vessel (backed by evidence from a computational fluid dynamics study (*16*)). A mean-flow approximation for the pulsatile flow in the umbilical artery(ies) is justified by the tissue diffusion that occurs over timescales much longer than a typical fetal heart cycle (*15*). The extent of coupling between the cord’s vessels depends on the cord helicity *Ω*^∗^ and the specific vascular configuration in the cord’s cross-section, including vessel-separation distance *d* and characteristic angles that describe the vessels’ positions within the cord (see Fig. 4 and the Supplement for more details). The total flux exchanged between the artery(ies) and the vein(s) per unit cord length is given by 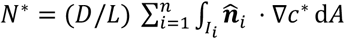, where *L* is the cord length, *D* is the tissue diffusivity, and 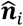 is the unit normal pointing outward each vessel’s interface *Ii* with the tissue (see Supplementary Table S3 for an illustration of cross-sectional multivessel fluxes).

Similar to Zwicker *et al*. (*54*), we define a metric of cord transfer efficiency 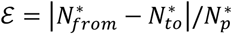 that quantifies the flux of solute or heat delivered to (or removed from) the fetus by the counter-current cord system relative to the placental exchange flux 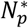 (see Supplement, Eq. S7). Unlike classical multistream heat-exchanger effectiveness (*55*), this physiological metric is fetus-centred, aiming to characterize the impact of cord shunting on net feto-placental transfer. For two-vessel cords,

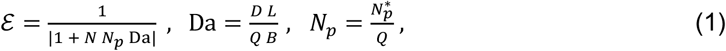

where Da is the dimensionless Damköhler number, a flux-ratio of the diffusive inter-vessel exchange and the advective transport along the cord (*Q* is the net umbilical flow rate, and *B* is the advective facilitation factor due to haemoglobin binding: *B* ∼ 100 for oxygen and *B* = 1 for heat; see Supplement for more details); *N* represents the scaled flux exchanged between the artery and the vein in a cross-sectional plane of the cord, which depends on specific cord geometry. Eq. (1) also holds for three-vessel cords with equipartitioned flow and placental supply between the two arteries. For four-vessel configurations, additional inter-vessel exchange fluxes in the cross-sectional plane were calculated numerically in COMSOL Multiphysics v. 6.0 (see Supplement Tables S1-S3).

## Acknowledgments

We gratefully acknowledge the Center for Perinatal Discovery at the University of California San Diego for making available Figs 1(d) and 4(c) from Professor Benirschke’s image library (Ref. (*45*)).

## Author Contributions

OEJ, RML and ILC designed research; TW and DL performed research and analysed data; SNS, NC, HKL, EDJ and RML contributed new data and analysis tools; TW, DL, OEJ and ILC wrote the first draft; all authors interpreted results and revised the paper.

## Data, Code, and Materials Availability

This study did not generate new materials. All data and code needed to evaluate and reproduce the results in the paper are present in the paper and/or the Supplementary Materials. The associated structural datasets and computational codes are deposited in Figshare repository: https://doi.org/10.6084/m9.figshare.30417331.

## References

1. M. Carter, Genomics, the diversification of mammals, and the evolution of placentation. Dev. Biol. 516, 167–182 (2024).

2. D. Laundon, N. J. Gostling, B. G. Sengers, P. Chavatte-Palmer, R. M. Lewis, Placental evolution from a three-dimensional and multiscale structural perspective. Evolution 78, 13–25 (2023).

3. Capellini, C. Venditti, R. A. Barton, Placentation and maternal investment in mammals. Am. Nat. 177, 86–98 (2011).

4. Capellini, C. L. Nunn, R. A. Barton, Microparasites and placental invasiveness in eutherian mammals. PLOS ONE 10, e0132563 (2015).

5. D. Laundon, B. G. Sengers, J. Thompson, S. E. Harris, O. Beasley, P. J. Basford, O. L. Katsamenis, P. Goggin, E. Derisoud, D. Fanelli, C. Bocci, F. Camillo, J. Shotton, G. Constable-Dakeyne, N. J. Gostling, P. Chavatte-Palmer, R. M. Lewis, Convergently evolved placental villi show multiscale structural adaptations to differential placental invasiveness. Biol. Lett. 20, 20240016 (2024).

6. D. Laundon, N. J. Gostling, I. G. Reddin, B. G. Sengers, P. Chavatte-Palmer, R. M. Lewis, Placental crises: disruptive selection and maternal under-investment as the foundations of mammalian placental evolution and dysfunction. Biol. Rev. 101, 1627–1640 (2026).

7. A. Erlich, P. Pearce, R. P. Mayo, O. E. Jensen, I. L. Chernyavsky, Physical and geometric determinants of transport in fetoplacental microvascular networks. Sci. Adv. 5, eaav6326 (2019).

8. N. Panitchob, K. L. Widdows, I. P. Crocker, E. D. Johnstone, C. P. Please, C. P. Sibley, J. D. Glazier, R. M. Lewis, B. G. Sengers, Computational modelling of placental amino acid transfer as an integrated system. Biochim. Biophys. Acta BBA - Biomembr. 1858, 1451–1461 (2016).

9. K. Benirschke, G. J. Burton, R. N. Baergen, “Anatomy and Pathology of the Umbilical Cord” in Pathology of the Human Placenta (Springer, Berlin, Heidelberg, ed. 6th, 2012; 10.1007/978-3-642-23941-0_12), pp. 309–375.

10. H. P. Laburn, D. Mitchell, K. Goelst, Fetal and maternal body temperatures measured by radiotelemetry in near-term sheep during thermal stress. J. Appl. Physiol. 72, 894–900 (1992).

11. G. G. Power, H. Schroder, R. D. Gilbert, Measurement of fetal heat production using differential calorimetry. J. Appl. Physiol. 57, 917–922 (1984).

12. G. B. West, W. H. Woodruff, J. H. Brown, Allometric scaling of metabolic rate from molecules and mitochondria to cells and mammals. Proc. Natl. Acad. Sci. 99, 2473–2478 (2002).

13. R. D. Gilbert, H. Schroder, T. Kawamura, P. S. Dale, G. G. Power, Heat transfer pathways between fetal lamb and ewe. J. Appl. Physiol. 59, 634–638 (1985).

14. H. Asakura, Fetal and neonatal thermoregulation. J. Nippon Med. Sch. 71, 360–370 (2004).

15. T. Wan, E. D. Johnstone, S. N. Saw, O. E. Jensen, I. L. Chernyavsky, A functional shunt in the umbilical cord: the role of coiling in solute and heat transfer. J. R. Soc. Interface 22, 20250148 (2025).

16. D. Kasiteropoulou, A. Topalidou, S. Downe, A computational fluid dynamics modelling of maternal-fetal heat exchange and blood flow in the umbilical cord. PLOS ONE 15, e0231997 (2020).

17. E. O. Janosko, J. Z. Jona, R. P. Belin, Congenital anomalies of the umbilicus. Am. Surg. 43, 177–185 (1977).

18. J. Spurway, P. Logan, S. Pak, The development, structure and blood flow within the umbilical cord with particular reference to the venous system. Australas. J. Ultrasound Med. 15, 97–102 (2012).

19. L. Murphy-Kaulbeck, L. Dodds, K. S. Joseph, M. Van den Hof, Single umbilical artery risk factors and pregnancy outcomes. Obstet. Gynecol. 116, 843–850 (2010).

20. S. Puvabanditsin, E. Garrow, M. Bhatt, S. Kathiravan, S. Gowda, R. Wong, M. Nagar, Four-vessel umbilical cord associated with multiple congenital anomalies: a case report and literature review. Fetal Pediatr. Pathol. 30, 98–105 (2011).

21. E. Jauniaux, C. Donner, P. Simon, M. Vanesse, J. Hustin, F. Rodesch, Pathologic aspects of the umbilical cord after percutaneous umbilical blood sampling. Obstet. Gynecol. 73, 215–218 (1989).

22. C. C. V. Dijk, A. Franx, M. W. M. de Laat, H. W. Bruinse, G. H. A. Visser, P. G. J. Nikkels, The umbilical coiling index in normal pregnancy. J. Matern. Fetal Neonatal Med. 11, 280–283 (2002).

23. G. A. Machin, J. Ackerman, E. Gilbert-Barness, Abnormal Umbilical Cord Coiling Is Associated with Adverse Perinatal Outcomes. Pediatr. Dev. Pathol. 3, 462–471 (2000).

24. C. Lees, G. Albaiges, C. Deane, M. Parra, K. H. Nicolaides, Assessment of umbilical arterial and venous flow using color Doppler. Ultrasound Obstet. Gynecol. 14, 250–255 (1999).

25. S. R. M. Reynolds, The proportion of Wharton’s jelly in the umbilical cord in relation to distention of the umbilical arteries and vein, with observations on the folds of Hoboken. Anat. Rec. 113, 365–377 (1952).

26. A. L. Fowden, P. M. Taylor, K. L. White, A. J. Forhead, Ontogenic and nutritionally induced changes in fetal metabolism in the horse. J. Physiol. 528, 209–219 (2000).

27. M. Carter, The blood supply to the abdominal organs of the fetal guinea-pig. J. Dev. Physiol. 6, 407–416 (1984).

28. J. P. Holt, E. A. Rhode, H. Kines, Ventricular volumes and body weight in mammals. Am. J. Physiol. 215, 704–715 (1968).

29. R. S. Seymour, Q. Hu, E. P. Snelling, C. R. White, Interspecific scaling of blood flow rates and arterial sizes in mammals. J. Exp. Biol. 222, jeb199554 (2019).

30. V. M. Savage, A. P. Allen, J. H. Brown, J. F. Gillooly, A. B. Herman, W. H. Woodruff, G. B. West, Scaling of number, size, and metabolic rate of cells with body size in mammals. Proc. Natl. Acad. Sci. 104, 4718–4723 (2007).

31. R. V. Lacro, K. L. Jones, K. Benirschke, The umbilical cord twist: origin direction, and relevance. Am. J. Obstet. Gynecol. 157, 833–838 (1987).

32. M. W. M. de Laat, P. G. J. Nikkels, A. Franx, G. H. A. Visser, The Roach muscle bundle and umbilical cord coiling. Early Hum. Dev. 83, 571–574 (2007).

33. P. Todtenhaupt, T. B. Kuipers, K. L. Dijkstra, L. M. Voortman, L. A. Franken, J. A. Spekman, T. H. Jonkman, S. G. Groene, A. A. Roest, M. C. Haak, Ej. T. Verweij, M. van Pel, E. Lopriore, B. T. Heijmans, L. E. van der Meeren, Twisting the theory on the origin of human umbilical cord coiling featuring monozygotic twins. Life Sci. Alliance 7, e202302543 (2024).

34. M. Vandeplassche, H. Lauwers, The twisted umbilical cord: an expression of kinesis of the equine fetus? Anim. Reprod. Sci. 10, 163–175 (1986).

35. M. E. Miller, M. Higginbottom, D. W. Smith, Short umbilical cord: its origin and relevance. Pediatrics 67, 618–621 (1981).

36. C. Moessinger, W. A. Blanc, P. A. Marone, D. C. Polsen, Umbilical cord length as an index of fetal activity: experimental study and clinical implications. Pediatr. Res. 16, 109–112 (1982).

37. K. C. Smith, A. S. Blunden, K. E. Whitwell, K. A. Dunn, A. D. Wales, A survey of equine abortion, stillbirth and neonatal death in the UK from 1988 to 1997. Equine Vet. J. 35, 496–501 (2003).

38. H. Asakura, K. T. Ball, G. G. Power, Interdependence of arterial PO2 and O2 consumption in the fetal sheep. J. Dev. Physiol. 13, 205–213 (1990).

39. H. O. Morishima, M. N. Yeh, W. H. Niemann, L. S. James, Temperature gradient between fetus and mother as an index for assessing intrauterine fetal condition. Am. J. Obstet. Gynecol. 129, 443–448 (1977).

40. J. C. K. Wells, Thermal environment and human birth weight. J. Theor. Biol. 214, 413–425 (2002).

41. P. Lakhoo, N. Brink, L. Radebe, M. H. Craig, M. D. Pham, M. M. Haghighi, A. Wise, I. Solarin, S. Luchters, G. Maimela, M. F. Chersich, A. Chikandiwa, B. Nakstad, C. Y. Wright, C. Brimicombe, K.-G. Technau, L. Harden, M. Boeckmann, R. Strehlau, R. S. Hetem, S. Luchters, G. Maimela, L. Radebe, M. Chersich, D. P. Lakhoo, I. Solarin, N. Brink, Heat-Health Study Group, HIGH Horizons Study Group, A systematic review and meta-analysis of heat exposure impacts on maternal, fetal and neonatal health. Nat. Med. 31, 684–694 (2025).

42. H. P. Laburn, How does the fetus cope with thermal challenges? Physiology 11, 96–100 (1996).

43. G. Pennati, K. Laganà, F. Gervaso, S. Rigano, E. Ferrazzi, How do cord compressions affect the umbilical venous flow resistance? An in vitro investigation of the biomechanical mechanisms. Cardiovasc. Eng. Technol. 4, 267–275 (2013).

44. S. N. Saw, Y. Dai, C. H. Yap, A review of biomechanics analysis of the umbilical–placenta system with regards to diseases. Front. Physiol. 12, 587635 (2021).

45. Benirschke, K., Comparative Placentation (2012). http://placentation.ucsd.edu.

46. Laundon, E. Proudley, P. J. Basford, O. L. Katsamenis, D. S. Chatelet, J. K. Cleal, N. J. Gostling, P. Chavatte-Palmer, R. M. Lewis, Quantitative microCT imaging of a whole equine placenta and its blood vessel network. Placenta 154, 216–219 (2024).

47. Belevich, M. Joensuu, D. Kumar, H. Vihinen, E. Jokitalo, Microscopy image browser: a platform for segmentation and analysis of multidimensional datasets. PLOS Biol. 14, e1002340 (2016).

48. L. J. Revell, phytools 2.0: an updated R ecosystem for phylogenetic comparative methods (and other things). PeerJ 12, e16505 (2024).

49. Paradis, K. Schliep, ape 5.0: an environment for modern phylogenetics and evolutionary analyses in R. Bioinformatics 35, 526–528 (2019).

50. N. S. Upham, J. A. Esselstyn, W. Jetz, Inferring the mammal tree: species-level sets of phylogenies for questions in ecology, evolution, and conservation. PLOS Biol. 17, e3000494 (2019).

51. W. Jetz, R. Guralnick, R. Bowie, A. Pyron, J. Esselstyn, VertLife: Vertebrate taxa, mammals (2025). https://vertlife.org/data/mammals/.

52. P. O. Lewis, A likelihood approach to estimating phylogeny from discrete morphological character data. Syst. Biol. 50, 913–925 (2001).

53. P. Virtanen, R. Gommers, T. E. Oliphant, M. Haberland, T. Reddy, D. Cournapeau, E. Burovski, P. Peterson, W. Weckesser, J. Bright, S. J. van der Walt, M. Brett, J. Wilson, K. J. Millman, N. Mayorov, A. R. J. Nelson, E. Jones, R. Kern, E. Larson, C. J. Carey, I. Polat, Y. Feng, E. W. Moore, J. VanderPlas, D. Laxalde, J. Perktold, R. Cimrman, I. Henriksen, E. A. Quintero, C. R. Harris, A. M. Archibald, A. N. H. Ribeiro, F. Pedregosa, P. van Mulbregt, SciPy 1.0 Contributors, SciPy 1.0: fundamental algorithms for scientific computing in Python. Nat. Methods 17, 261–272 (2020).

54. D. Zwicker, R. Ostilla-Mónico, D. E. Lieberman, M. P. Brenner, Physical and geometric constraints shape the labyrinth-like nasal cavity. Proc. Natl. Acad. Sci. 115, 2936–2941 (2018).

55. D. P. Sekulić, R. K. Shah, “Thermal design theory of three-fluid heat exchangers” in Advances in Heat Transfer (Elsevier, 1995), vol. 26, pp. 219–328.

56. M. V. Blanco, H. R. Vega, R. A. Guerri-Guttenberg, R. Giuliano, D. R. Grana, F. Azzato, J. Milei, Histopathology and histomorphometry of umbilical cord blood vessels. Findings in normal and high risk pregnancies. Artery Res. 5, 50–57 (2011).

57. Pierre, J. Bouyssier, F. de Gournay, F. Plouraboué, Numerical computation of 3D heat transfer in complex parallel heat exchangers using generalized Graetz modes. J. Comput. Phys. 268, 84–105 (2014).

58. W. Käsinger, R. Huch, A. Huch, In vivo oxygen dissociation curve for whole fetal blood: fitting the Adair equation and blood gas nomogram. Scand. J. Clin. Lab. Invest. 41, 701–707 (1981).

59. D. Aulds, R. F. Barron, Three-fluid heat exchanger effectiveness. Int. J. Heat Mass Transf. 10, 1457–1462 (1967).

60. Shrivastava, T. A. Ameel, Three-fluid heat exchangers with three thermal communications. Part B: effectiveness evaluation. Int. J. Heat Mass Transf. 47, 3867–3875 (2004).

61. D.-F. Ruan, X.-F. Yuan, S.-Y. Wu, Y.-R. Li, Exergy effectiveness analysis of three-fluid heat exchanger. J. Supercond. Nov. Magn. 23, 1127–1131 (2010).

62. O. E. Jensen, I. L. Chernyavsky, Blood flow and transport in the human placenta. Annu. Rev. Fluid Mech. 51, 25–47 (2019).

63. R. Lee, T. M. Mayhew, Star volumes of villi and intervillous pores in placentae from low and high altitude pregnancies. J. Anat. 186, 349–355 (1995).

64. A. Weissman, P. Jakobi, M. Bronshtein, I. Goldstein, Sonographic measurements of the umbilical cord and vessels during normal pregnancies. J. Ultrasound Med. Off. J. Am. Inst. Ultrasound Med. 13, 11–14 (1994).

65. D. Kaplan, A. J. Jaffa, I. E. Timor, D. Elad, Hemodynamic analysis of arterial blood flow in the coiled umbilical cord. Reprod. Sci. Thousand Oaks Calif 17, 258–268 (2009).

66. M. de Laat, A. Franx, E. van Alderen, P. Nikkels, G. Visser, The umbilical coiling index, a review of the literature. J. Matern.-Fetal Neonatal Med. Off. J. Eur. Assoc. Perinat. Med. Fed. Asia Ocean. Perinat. Soc. Int. Soc. Perinat. Obstet. 17, 93–100 (2005).

67. J. Jordan, E. Ackerman, R. L. Berger, Polarographic diffusion coefficients of oxygen defined by activity gradients in viscous media. J. Am. Chem. Soc. 78, 2979–2983 (1956).

68. Ponder, The coefficient of thermal conductivity of blood and of various tissues. J. Gen. Physiol. 45, 545–551 (1962).

69. A. Nye, E. Ingram, E. D. Johnstone, O. E. Jensen, H. Schneider, R. M. Lewis, I. L. Chernyavsky, P. Brownbill, Human placental oxygenation in late gestation: experimental and theoretical approaches. J. Physiol. 596, 5523–5534 (2018).

70. D. Walker, A. Walker, C. Wood, Temperature of the human fetus. J. Obstet. Gynaecol. J. Inst. Obstet. Gynaecol. 76, 503–511 (1969).

