## Supplementary Materials for "Umbilical cord structure shapes feto-maternal heat transfer across mammals"

**This PDF file includes:**

Supplementary Materials and Methods

Figures S1 to S4

Tables S1 to S3

This Supplement provides further details on the governing equations and computational methodology (Sec. S1-S2), parameter estimation (Sec. S3-S4), and additional results (Figs S3-S4). The associated structural datasets and computational codes are publicly available in the Figshare repository: `https://doi.org/10.6084/m9.figshare.30417331`.

### Supplementary Materials and Methods

#### S1. A three-dimensional model of solute exchange between helical vessels

We build a three-dimensional theoretical model of solute exchange between helical vessels in an umbilical cord. We solve the steady diffusion equation in the umbilical cord accounting for its helical structure and calculate the total solute flux between vessels. The model rests on the following assumptions. The cord is straight and vessel centerlines form helical curves with a common axis running along the centre of the cord; all helices have the same uniform pitch $2\pi/\Omega^*$; vessel boundaries are circles in any plane orthogonal to the cord axis, and vessel boundaries in any such plane are identical up to rotation; the pitch $2\pi/\Omega^*$ is comparable in magnitude to the cord radius $R_c^*$, allowing for significant twisting; the cord length $L^*$ is large compared to $R_c^*$; secondary flows ensure that concentration profiles of solutes within each vessel are uniform within any cross-section, but vary slowly over the length of the cord; temporal fluctuations are ignored. We then develop an approximation exploiting the relative magnitudes of the two geometric parameters $\Omega = \Omega^* R_c^* = O(1)$ and $\epsilon = R_c^*/L^* \ll 1$. In any cross-sectional plane, the solute diffuses between vessels at a rate influenced by $\Omega$; here we use an existing model [15] to calculate the relevant fluxes, exploiting the geometric self-similarity in vessel profiles along the cord. Along the length of the cord, we model slow axial changes in vessel concentrations arising from inter-vessel fluxes, and introduce a solute flux from the placenta at one end of the cord. This allows us to evaluate the impact of quasi-steady counter-current exchange within the cord on solute transport from placenta to fetus, accounting for vessel twisting.

**Vessel geometry.** We model the umbilical cord as containing $n + m$ helical blood vessels ($n$ arteries and $m$ veins, numbering arteries from $i = 1$ to $n$ and veins from $i = n + 1$ to $i = n + m$). Let $\mathbf{x}^* = (x^*, y^*, z^*)$ be a dimensional Cartesian coordinate system such that the unit vector $\hat{\mathbf{z}}$ lies along the axis of the cord. We assume that the centre of one vein ($i = n + 1$) intersects $(R_v^*, 0, 0)$ where $R_v^*$ is the vein's helical radius (i.e., the vein centreline lies on a cylinder of radius $R_v^*$).

Introducing the variable $s$, the vein's centreline lies along

$$x^* = R_v^* \cos s, \quad y^* = R_v^* \sin s, \quad z^* = s/\Omega^* \quad \text{for} \quad s \in [0, \Omega^* L^*]; \tag{S1}$$

$s = 0$ at the fetal end of the cord and $s = \Omega^* L^*$ at the maternal end of the cord. This reference vein is chosen such that any other veins lie in the third or fourth quadrant of the cross-sectional plane (see Fig. S1a). We assume that all arteries have same cross-sectional radius $r_a^*$ and all veins have same cross-sectional radius $r_v^*$, taking $R_v^* + r_v^* < R_c^*$ to ensure the reference vein lies fully within the cord. The distance between the centre of the reference vein and the centre of any artery is $R_{va}^* \equiv d_a^* + r_v^* + r_a^*$, where $d_a^* > 0$ is the minimum distance between any artery and the reference vein. We assume $d_a^* + 2r_a^* + 2r_v^* < R_c^*$, so that arteries lie entirely within the cord. Similarly, we define $R_{vv}^* \equiv d_v^* + 2r_v^*$ to be the in-plane distance between the centre of the reference vein and the centre of any other vein, where $d_v^* > 0$ is the minimum separation between vein $i$ $(i = n+2, ..., n+m)$ and the reference vein, and $d_v^* + 4r_v^* < R_c^*$. We define $\theta_i$ for $i = 1, ..., n$ and $i = n+2, \ldots, n+m$ to be the angle subtended by the centre of vessel $i$ with the $x$-axis at the centre of the reference vein in the plane $z^* = 0$, and ensure differences between any two $\theta_i$ are large enough to avoid vessels intersecting. Each vessel is built by sweeping the centre of a circle along the centreline of a helix, so that vessel centrelines lie along

$$x_i^*(s) = R_v^* \cos s + R_{va}^* \cos(s + \theta_i), \quad y_i^*(s) = R_v^* \sin s + R_{va}^* \sin(s + \theta_i), \quad (i = 1, \ldots, n), \tag{S2a}$$

$$x_i^*(s) = R_v^* \cos s + R_{vv}^* \cos(s + \theta_i), \quad y_i^*(s) = R_v^* \sin s + R_{vv}^* \sin(s + \theta_i), \quad (i = n+2, \ldots, n+m), \tag{S2b}$$

with $z^* = s/\Omega^*$, $0 \leq s \leq \Omega^* L^*$. The helical radius $R_i^*$ of vessel $i$ is the distance between the centre of the cord and the centre of vessel $i$. Thus

$$R_i^* = \begin{cases} \sqrt{(R_{va}^*)^2 + (R_v^*)^2 + 2R_v^* R_{va}^* \cos(\theta_i)} & (i = 1, \ldots, n), \\ R_v^* & (i = n+1), \\ \sqrt{(R_{vv}^*)^2 + (R_v^*)^2 + 2R_v^* R_{vv}^* \cos(\theta_i)} & (i = n+2, \ldots, n+m). \end{cases} \tag{S3}$$

Vessel positions in the cross-section of a four-vessel cord (two arteries and two veins) in the plane $z^* = 0$ are illustrated in Fig. S1.

Let $I_0$ be the cylindrical domain occupied by the cord and vessels and $I_i$ for $i = 1, ..., n+m$ be the domain occupied by vessel $i$. We define $\partial I_i$ for $i = 0, 1, ..., n+m$ to be the corresponding vessel boundaries; the boundary $\partial I_0$ of the cord domain is shown in Fig. S2(a,b) for a four-vessel cord. The cord tissue volume is $I_t \equiv I_0 \setminus (\cup_{i=1}^{n+m} I_i)$. Denote a plane $\mathcal{P}$ where $z^*$ is constant. Then the line boundaries for the cord and each vessel on the $(x^*, y^*)$ plane for fixed $z^*$ can be written as $\partial S_i \equiv \partial I_i \cap \mathcal{P}$ for $i = 0, 1, ..., n+m$ respectively. The cord tissue surface in a plane for fixed $z^*$ is $S_t = I_t \cap \mathcal{P} = S_0 \setminus (\cup_{i=1}^{n+m} S_i)$, and we denote the end plates of the cord by $I_f \equiv I_t \cap \mathcal{P}\big|_{z^*=0}$ and $I_p \equiv I_t \cap \mathcal{P}\big|_{z^*=L^*}$ respectively (see Fig. S2c).

**Solute transport.** Dimensional solute concentration (or temperature) $c^*(x^*, y^*, z^*)$ in the cord tissue $I_t$ is modelled using the steady diffusion equation

$$\nabla^2 c^* = 0 \quad \text{in } I_t, \tag{S4a}$$

$$\hat{\mathbf{n}} \cdot \nabla c^* = 0 \quad \text{on } \partial I_0,\ I_f \text{ and } I_p, \tag{S4b}$$

$$c^* = c_i^*(z^*) \quad \text{on } \partial I_i, \text{ for } i = 1, ..., n+m. \tag{S4c}$$

No-flux conditions (S4b) are imposed on the cord's outer boundary $\partial I_0$, the cross-sections of the umbilical cord at the fetal end $I_f$ and the placental end $I_p$, with $\hat{\mathbf{n}}$ denoting a unit normal pointing out of the tissue. In (S4c), we prescribe axially varying solute concentration $c_i^*(z^*)$ on the surface of vessel $i$ ($i = 1, \ldots, n+m$). Secondary flows induced by the helical geometry promote the mixing of solutes in the vessels, partially supporting the assumption that the concentration is uniform across any plane $z^* =$ constant. However, for straight vessels, the impact of solute heterogeneity may not be negligible [57]. Boundary condition (S4c) treats flow-to-wall heat transfer within each vessel as fast relative to tissue diffusion; finite thermal inertia or azimuthal non-uniformity of the wall would reduce absolute exchange coefficients and may be most relevant for very closely-spaced vessels or straight cords. However, as indicated by three-dimensional fluid dynamics simulations [16], these effects should not affect the qualitative geometry-dependent trends considered in this study.

Solving (S4) allows us in principle to evaluate the solute flux per unit length $D^* \hat{J}_i^*(z^*)$ entering vessel $i$. The cross-sectionally-averaged transport equation in each vessel can then be written

$$BQ_i^* \frac{\mathrm{d}}{\mathrm{d}z^*} c_i^*(z^*) = D^* \hat{J}_i^*(z^*) \quad \text{for } i = 1, ..., n, \tag{S5a}$$

$$-BQ_i^* \frac{\mathrm{d}}{\mathrm{d}z^*} c_i^*(z^*) = D^* \hat{J}_i^*(z^*) \quad \text{for } i = n+1, ..., n+m. \tag{S5b}$$

When modelling heat transport, $B = 1$. When modelling oxygen transport, the parameter $B$ describes the linearised oxygen-haemoglobin binding kinetics in the fetal blood that boosts its advective capacity [58]. $D^*$ is the diffusion coefficient in tissue and $Q_i^*$ for $i = 1, ..., n+m$ represents axial blood volume flux within each vessel. We assume that the total umbilical volumetric blood flow rate $Q^*$ is conserved between arteries and veins, so that $Q^* = \sum_{i=1}^{n} Q_i^* = \sum_{j=n+1}^{n+m} Q_j^*$. The negative sign in (S5b) arises because the flow in the veins travels in the opposite direction to the flow in the arteries (see Fig. S2d). For vessels separated by a distance $d^*$ within the cord, the diffusion time for oxygen $((d^*)^2/D^*$ is of magnitude 10 to 100s, which is long compared to the fetal heartbeat period (of order $10^{-1}$ s). We therefore ignore the rapid fluctuations of arterial flow.

We impose the following boundary conditions at the cord's fetal end $z^* = 0$ and placental end $z^* = L^*$:

$$c_i^*(0) = c_F^* \qquad (i = 1, \ldots, n), \tag{S6a}$$

$$c_i^*(L^*) = \frac{1}{mQ_i^*}\left[\sum_{j=1}^{n} Q_j^* c_j^*(L^*) + N_p^*\left(c_M^* - \frac{1}{n}\sum_{j=1}^{n} c_j^*(L^*)\right)\right] \qquad (i = n+1, \ldots, n+m). \tag{S6b}$$

Here, $c_F^*$ is the incoming fetal solute concentration. In (S6b) we include a flux (proportional to the factor $N_p^*$, defined in Section S3 below) modelling the placenta which depends on the difference between the average fetal arterial concentration at $z^* = L^*$ and the average maternal placental intervillous solute concentration $c_M^*$. This approximate boundary condition is based on the assumption that the arteries split into the placenta with equal weights; likewise we assume that the volumetric and solute fluxes leaving the placenta are uniformly distributed between veins.

We define a metric to describe the cord transfer efficiency, namely

$$\mathcal{E} = \frac{1}{N_p^*}\left|\frac{\sum_{i=n+1}^{n+m} Q_i^* c_i^*(0) - Q^* c_F^*}{c_M^* - c_F^*}\right|. \tag{S7}$$

$\mathcal{E} = 1$ is obtained when the solute flux delivered to the fetus, $\sum_{i=n+1}^{n+m} Q_i^* c_i^*(0)$ is equal to the sum of the solute flux supplied by the placenta and the inlet flux, $N_p^*(c_M^* - c_F^*) + Q^* c_F^*$. $\mathcal{E} = 0$ when the solute flux delivered to the fetus $\sum_{i=n+1}^{n+m} Q_i^* c_i^*(0)$ is the same as the inlet flux $Q^* c_F^*$.

**Nondimensionalisation.** We scale parameters and variables using $R_c^*$ as a reference length and obtain the dimensionless geometric parameters (without asterisks) shown in Table S1. For simplicity we assume $d_a = d_v = d$, say. Among these parameters, we highlight $\Omega = \Omega^* R_c^* = O(1)$ and $\epsilon = R_c^*/L^* \equiv 1/L \ll 1$. Typical values of physico-chemical parameters are given in Table S2. From these we construct a Damköhler number

$$\mathrm{Da} = \frac{D^* L^*}{Q^* B} \tag{S8}$$

that measures the ratio between diffusive and advective transport along the cord; its magnitude depends on the type of solute considered (via $D^*$ and $B$) and mammal characteristics (via $Q^*$ and $L^*$).

We scale concentrations relative to fixed maternal and fetal arterial concentrations $c_M^*$ and $c_F^*$ as

$$c^*(x^*, y^*, z^*) = c_M^* + (c_F^* - c_M^*)\, c(x, y, z), \quad c_i^*(z^*) = c_M^* + (c_F^* - c_M^*)\, c_i(z) \tag{S9}$$

and solute fluxes such that $\hat{J}_i^* = (c_F^* - c_M^*)\hat{J}_i$ $(i = 1, ..., n + m)$. For oxygen, $c_F^* - c_M^* < 0$ since maternal oxygen concentration is significantly higher than fetal concentration, while for heat $c_F^* - c_M^* > 0$ since fetal temperature is higher than maternal temperature. We also define dimensionless fluxes

$$q_i = \frac{Q_i^*}{Q^*}, \text{ for } i = 1, ..., n + m \text{ where } \sum_{i=1}^{n} q_i = \sum_{j=n+1}^{n+m} q_j = 1, \tag{S10}$$

$$N_p^* = Q^* N_p. \tag{S11}$$

Here $q_i$ represents the proportion of the total umbilical blood flow $Q^*$ that passes through vessel $i$ and $N_p$ is the placental heat or solute flux scaled with the flux carried by umbilical blood flow.

Expressed in nondimensional variables, (S4) becomes

$$\nabla^2 c = 0 \quad \text{in } I_t, \tag{S12a}$$

$$\hat{\mathbf{n}}_0 \cdot \nabla c = 0 \quad \text{on} \quad \partial I_0, \tag{S12b}$$

$$\hat{\mathbf{n}} \cdot \nabla c = 0 \quad \text{on } S_0 \text{ and on } S_L, \tag{S12c}$$

$$c = c_i(z) \text{ on } \partial I_i \quad \text{for } i = 1, ..., n + m. \tag{S12d}$$

The flux entering vessel $i$ through its curved boundaries satisfies

$$\int_0^{1/\epsilon} \hat{J}_i \,\mathrm{d}z = \int_{\partial I_i} \hat{\mathbf{n}}_i \cdot \nabla c \,\mathrm{d}A \qquad (i = 1, \ldots, n + m). \tag{S13}$$

On the cord boundary $\partial I_0$, $\hat{\mathbf{n}}_0 = (x, y, 0)$ on $x^2 + y^2 = 1$. The other normal vectors, $\hat{\mathbf{n}}_i$ for $i = 1, ..., n + m$, typically have a component that lies outside any plane $z =$ constant (see Fig. S2a for a four-vessel cord example). The scaled transport equations (S5) take the form

$$q_i \partial_z c_i(z) = \epsilon \mathrm{Da} \hat{J}_i \quad \text{for } i = 1, ..., n, \tag{S14a}$$

$$-q_j \partial_z c_j(z) = \epsilon \mathrm{Da} \hat{J}_j \quad \text{for } j = n + 1, ..., n + m, \tag{S14b}$$

and the boundary conditions (S6) become

$$c_i = 1 \text{ for } i = 1, ..., n \quad \text{at } z = 0, \tag{S15a}$$

$$c_j = \frac{1}{mq_j} \left[ \sum_{i=1}^{n} q_i c_i - N_p \left( \frac{1}{n} \sum_{i=1}^{n} c_i \right) \right] \text{ for } j = n + 1, ..., n + m \quad \text{at } z = L. \tag{S15b}$$

The cord efficiency (S7) becomes

$$\mathcal{E} = \frac{\left|1 - \sum_{j=n+1}^{n+m} q_j c_j(0)\right|}{N_p}. \tag{S16}$$

Our aim is to solve the countercurrent model (S14) accounting for the impact of helicity on the exchange between the vessels. For simplicity, motivated by the cord's primary physiological function, we focus on the loss of cord transfer efficiency associated with artery–vein exchange, rather than on broader multi-vessel exchange metrics [55, 59–61].

**Asymptotic model reduction.** In the limit $\epsilon \to 0$, keeping $\Omega$, Da and other geometric parameters of order unity, we see from (S14) that axial concentration gradients will be small, varying with the rescaled variable $\zeta = \epsilon z \equiv z/L$. Furthermore, we expect cross-vessel concentration profiles to share the self-similar rotation of the vessel profiles. Following [15] we therefore introduce a rotating coordinate system $\mathbf{X} = (X, Y, Z)$, where

$$X = x\cos(\Omega z) + y\sin(\Omega z), \quad Y = -x\sin(\Omega z) + y\cos(\Omega z), \quad Z = z, \tag{S17}$$

such that the vessels' (circular) cross-sections in any plane $\mathcal{P}$ lying perpendicular to the axis of the cord remain fixed with respect to $\mathbf{X}$. We then write the tissue and vessel solute concentrations as

$$c(\mathbf{x}) = C^{(0)}(\mathbf{X}_\perp, \zeta) + O(\epsilon), \qquad c_i(z) = C_i(\zeta). \tag{S18}$$

where $\mathbf{X}_\perp \equiv (X, Y)$. Because axial concentration gradients are weak, we recover from (S12), at leading order in $\epsilon$, the diffusion problem addressed in [15], namely

$$\nabla_\perp \circ \left(\nabla_\perp C^{(0)} + \Omega^2 \mathbf{H}^{(0)}\right) = 0 \quad \text{on } S_t, \tag{S19a}$$

$$\hat{\mathbf{m}}_0 \circ \nabla_\perp C^{(0)} = 0 \quad \text{on} \quad \partial S_0, \tag{S19b}$$

$$C^{(0)} = C_i(\zeta) \text{ on } \partial S_i, \quad \text{for } i = 1, ..., n+m, \tag{S19c}$$

where $\mathbf{H}^{(0)} \equiv \mathbf{X}^\perp(\mathbf{X}^\perp \circ \nabla_\perp C^{(0)})$. Here $\nabla_\perp \equiv (\partial_X, \partial_Y, 0)$, $\mathbf{X}^\perp \equiv (Y, -X, 0)$, $\hat{\mathbf{m}}_0 \equiv (X, Y, 0)$ on $X^2 + Y^2 = 1$ and the product $\circ$ is defined as $(a, b, c) \circ (d, e, f) \equiv ad + be + cf$. In general, this is not equivalent to a scalar product since the helical coordinate system $(X, Y, Z)$ is not orthogonal. The term proportional to $\Omega^2$ shows how in-plane azimuthal concentration gradients drive azimuthal fluxes that are amplified at increasing radial distance from the centre of the cord. The solute flux per unit length across the boundary of vessel is

$$\hat{J}_i = \int_{\partial S_i} \hat{\mathbf{m}}_i \circ \left[\nabla_\perp C^{(0)} + \Omega^2 \mathbf{H}^{(0)}\right] \mathrm{d}s \quad \text{for } i = 0, .., n+m. \tag{S20}$$

where $\hat{\mathbf{m}}_i$ are the in-plane unit normals oriented out of boundaries $\partial S_i$. Transport in vessels satisfies

$$q_i \partial_\zeta C_i = \mathrm{Da} \hat{J}_i \quad \text{for } i = 1, ..., n \tag{S21a}$$

$$-q_j \partial_\zeta C_j = \mathrm{Da} \hat{J}_j \quad \text{for } j = n+1, ..., n+m, \tag{S21b}$$

and the boundary conditions (S6) become

$$C_i = 1 \text{ for } i = 1, ..., n \quad \text{at } \zeta = 0, \tag{S22a}$$

$$C_j = \frac{1}{m q_j} \left[ \sum_{i=1}^{n} q_i C_i - N_p \left( \frac{1}{n} \sum_{i=1}^{n} C_i \right) \right] \text{ for } j = n+1, ..., n+m \quad \text{at } \zeta = 1. \tag{S22b}$$

**Exploiting linearity.** The solute concentration in any cross-section, governed by (S19), involves transport between multiple vessels. It is convenient to exploit linearity, identifying exchange from individual vessels, by writing $C^{(0)}$ as the combination

$$C^{(0)}(\mathbf{X}_\perp, \zeta) = \sum_{i=1}^{n+m} \psi_i(\mathbf{X}_\perp) C_i(\zeta), \tag{S23}$$

where each function $\psi_i$ for $i, j = 1, ..., n+m$ satisfies the canonical problem

$$\nabla_\perp \cdot \Big[ \nabla_\perp \psi_i + \Omega^2 \mathbf{X}^\perp (\mathbf{X}^\perp \cdot \nabla_\perp \psi_i) \Big] = 0, \tag{S24a}$$

$$\hat{\mathbf{m}}_0 \cdot \nabla_\perp \psi_i = 0 \quad \text{on} \quad \partial S_0, \tag{S24b}$$

$$\psi_i \big|_{\partial S_j} = \delta_{ij} \quad \text{where} \quad \delta_{ij} = \begin{cases} 1 & \text{if } i = j, \\ 0 & \text{if } i \neq j. \end{cases} \tag{S24c}$$

The boundary conditions (S24c) imply that $\sum_{i=1}^{n+m} \psi_i = 1$. Then, using (S23) to rewrite (S20) in terms of canonical solutions, the transport equations (S14) become

$$\pm q_i \, \partial_\zeta C_i(\zeta) = \mathrm{Da} \sum_{j=1}^{n+m} N_{ij}(\Omega) C_j(\zeta) \tag{S25}$$

(taking signs consistent with (S14)), where $N_{ij}(\Omega)$ is the flux for vessel $i$ for problem $\psi_j$ from (S24), defined as

$$N_{ij} = \int_{\partial S_i} \Big[ \hat{\mathbf{m}}_i \cdot \nabla_\perp \psi_j + \Omega^2 (\hat{\mathbf{m}}_i \cdot \mathbf{X}^\perp)(\mathbf{X}^\perp \cdot \nabla_\perp \psi_j) \Big] \, \mathrm{d}s. \tag{S26}$$

Writing $\mathbf{s}_i \equiv \nabla_\perp \psi_i + \Omega^2 \mathbf{X}^\perp (\mathbf{X}^\perp \cdot \nabla_\perp \psi_i)$ and applying the divergence theorem to $\nabla_\perp \cdot (\psi_i \mathbf{s}_j - \psi_j \mathbf{s}_i)$ reveals that $N_{ij} = N_{ji}$. Likewise applying the divergence theorem to $\nabla_\perp \cdot \mathbf{s}_j$ shows that $\sum_i N_{ij} = 0$.

Let $\mathbf{C} = (C_1, \ldots, C_n, C_{n+1}, \ldots, C_{n+m})^T$ and define matrices

$$\mathsf{Q} = \mathrm{diag}(q_1, \ldots, q_n, -q_{n+1}, \ldots, -q_{n+m}) \quad \text{and} \quad \{\mathsf{N}\}_{ij} = N_{ij}. \tag{S27}$$

We can then write the transport equations (S25) and boundary conditions (S22) as

$$\mathsf{Q}\,\partial_\zeta \mathbf{C} = \mathrm{Da}\,\mathsf{N}\mathbf{C}, \quad \begin{pmatrix} \mathsf{I}_n & 0 \\ 0 & 0 \end{pmatrix} \mathbf{C}(0) = \begin{pmatrix} 1_n \\ 0 \end{pmatrix}, \quad -m \begin{pmatrix} 0 & 0 \\ 0 & \mathsf{I}_m \end{pmatrix} \mathsf{Q}\mathbf{C}(1) = \begin{pmatrix} 0 \\ 1_m \end{pmatrix} \begin{pmatrix} 1_n \\ 0 \end{pmatrix}^T \Big(\mathsf{Q} - \tfrac{N_p}{n}\mathsf{I}\Big)\mathbf{C}(1). \tag{S28}$$

The problem (S28) for a cord configuration with an arbitrary number of vessels was solved with `Python`'s `SciPy` library. The function `scipy.integrate.solve_bvp` implements a collocation algorithm that iteratively approximates the solution of the differential equations by dividing the domain (in this case $\zeta \in [0, 1]$) into a set of discrete points and ensuring that boundary conditions are satisfied. This algorithm was used to obtain a numerical solution for a 4-vessel cord configuration. To obtain values for $\mathsf{N}$ in (S28), the minimum flux configuration in Fig. 4(f) was used (see Table S3).

### S2. Cord transfer efficiency calculation

When the cord contains two arteries and one vein ($n = 2$ and $m = 1$) we can derive an explicit solution to (S28). Using flow conservation between arteries and vein, and the symmetry $N_{ij} = N_{ji}$, we can define the parameters

$$q = \frac{Q_1^*}{Q^*}, \qquad 1 - q = \frac{Q_2^*}{Q^*}, \qquad \mathrm{Da}_{13} = \mathrm{Da}N_{13}, \qquad \mathrm{Da}_{23} = \mathrm{Da}N_{23}, \qquad \mathrm{Da}_{12} = \mathrm{Da}N_{12}. \tag{S29}$$

We can rewrite the transport equations as

$$q\partial_\zeta C_1 = \mathrm{Da}_{13}(C_3 - C_1) + \mathrm{Da}_{12}(C_2 - C_1), \tag{S30a}$$

$$(1-q)\partial_\zeta C_2 = \mathrm{Da}_{23}(C_3 - C_2) + \mathrm{Da}_{12}(C_1 - C_2), \tag{S30b}$$

$$\partial_\zeta C_3 = \mathrm{Da}_{13}(C_3 - C_1) + \mathrm{Da}_{23}(C_3 - C_2), \tag{S30c}$$

with boundary conditions

$$C_1 = C_2 = 1 \quad \text{at } \zeta = 0, \tag{S31a}$$

$$C_3 = qC_1 + (1-q)C_2 - \tfrac{1}{2}N_p(C_1 + C_2) \quad \text{at } \zeta = 1. \tag{S31b}$$

(S30) implies that

$$[C_3 - qC_1 - (1-q)C_2]_\zeta = 0, \tag{S32}$$

showing that the difference in concentration between the vein and the arteries is constant along the length of the cord. Integrating (S32) yields an expression for the concentration of the vein, $C_3$, in terms of the concentration of the two arteries and $K$, which represents the constant flux delivered by the placenta

$$C_3 = qC_1 + (1-q)C_2 + K \quad \text{for } 0 \le z \le 1. \tag{S33}$$

The constant $K = -\frac{1}{2}N_p(C_1(1) + C_2(1))$ is determined by applying (S31). Substituting (S33) in (S30b) and (S30c) gives

$$q\partial_\zeta C_1 = [\mathrm{Da}_{13}(1-q) + \mathrm{Da}_{12}]\,(C_2 - C_1) + K\mathrm{Da}_{13}, \tag{S34a}$$

$$(1-q)\partial_\zeta C_2 = [\mathrm{Da}_{23}q + \mathrm{Da}_{12}]\,(C_1 - C_2) + K\mathrm{Da}_{23}. \tag{S34b}$$

In matrix form, this is

$$\begin{pmatrix} C_1 \\ C_2 \end{pmatrix}_\zeta = \begin{pmatrix} -\gamma_1 & \gamma_1 \\ \gamma_2 & -\gamma_2 \end{pmatrix} \begin{pmatrix} C_1 \\ C_2 \end{pmatrix} + K \begin{pmatrix} \mu_1 \\ \mu_2 \end{pmatrix}, \tag{S35}$$

where

$$\gamma_1 = \frac{\mathrm{Da}_{13}(1-q) + \mathrm{Da}_{12}}{q}, \quad \gamma_2 = \frac{\mathrm{Da}_{23}q + \mathrm{Da}_{12}}{1-q}, \quad \mu_1 = \frac{\mathrm{Da}_{13}}{q} \quad \text{and} \quad \mu_2 = \frac{\mathrm{Da}_{23}}{1-q}. \tag{S36}$$

The full solution to this system can be written as the sum of the solution to the homogeneous problem and the solution to the particular integral,

$$\mathbf{C} = a\begin{pmatrix} 1 \\ 1 \end{pmatrix} + b\begin{pmatrix} \gamma_1 \\ -\gamma_2 \end{pmatrix} e^{-(\gamma_1+\gamma_2)\zeta}, \qquad \mathbf{C} = g(\zeta)\begin{pmatrix} 1 \\ 1 \end{pmatrix} + h(\zeta)\begin{pmatrix} \gamma_1 \\ -\gamma_2 \end{pmatrix}, \tag{S37}$$

for some constants $a, b$ and functions $g(\zeta), h(\zeta)$. Then, substituting (S37b) into (S35), we have

$$g_\zeta\begin{pmatrix} 1 \\ 1 \end{pmatrix} + h_\zeta\begin{pmatrix} \gamma_1 \\ -\gamma_2 \end{pmatrix} = -h\begin{pmatrix} \gamma_1^2 + \gamma_1\gamma_2 \\ -\gamma_1\gamma_2 - \gamma_2^2 \end{pmatrix} + K\begin{pmatrix} \mu_1 \\ \mu_2 \end{pmatrix}, \tag{S38}$$

so that

$$g_\zeta\begin{pmatrix} 1 \\ 1 \end{pmatrix} + [h_\zeta + (\gamma_1+\gamma_2)h]\begin{pmatrix} \gamma_1 \\ -\gamma_2 \end{pmatrix} = K\begin{pmatrix} \mu_1 \\ \mu_2 \end{pmatrix}. \tag{S39}$$

Multiply by $(1, -1)$ and $(\gamma_2, \gamma_1)$ to project onto the eigenvectors

$$[h_\zeta + (\gamma_1+\gamma_2)h](1,-1)\begin{pmatrix} \gamma_1 \\ -\gamma_2 \end{pmatrix} = K(1,-1)\begin{pmatrix} \mu_1 \\ \mu_2 \end{pmatrix}, \quad g_\zeta(\gamma_2,\gamma_1)\begin{pmatrix} 1 \\ 1 \end{pmatrix} = K(\gamma_2,\gamma_1)\begin{pmatrix} \mu_1 \\ \mu_2 \end{pmatrix}. \tag{S40}$$

Hence

$$g_\zeta = \frac{K(\mu_1\gamma_2 + \mu_2\gamma_1)}{\gamma_1+\gamma_2} \quad \text{and} \quad h_\zeta + (\gamma_1+\gamma_2)h = \frac{K(\mu_1 - \mu_2)}{\gamma_1+\gamma_2}. \tag{S41}$$

Then

$$g = K\left(\frac{\mu_1\gamma_2 + \mu_2\gamma_1}{\gamma_1+\gamma_2}\right)\zeta \quad \text{and} \quad h = \frac{K(\mu_1-\mu_2)}{(\gamma_1+\gamma_2)^2} \tag{S42}$$

are particular integrals, independent of the complementary functions in (S37). The full solution is the sum of the homogeneous solution and the particular integral in (S37), giving

$$\begin{aligned}\mathbf{C}(\zeta) = \begin{pmatrix} C_1 \\ C_2 \end{pmatrix} &= \begin{pmatrix} 1 \\ 1 \end{pmatrix}\left[1 - N_p\,\zeta\,\frac{C_1(1)+C_2(1)}{2}\left(\frac{\mu_1\gamma_2+\mu_2\gamma_1}{\gamma_1+\gamma_2}\right)\right] \\ &- \begin{pmatrix} \gamma_1 \\ -\gamma_2 \end{pmatrix}\frac{N_p(\mu_1-\mu_2)}{(\gamma_1+\gamma_2)^2}\frac{C_1(1)+C_2(1)}{2}\left[1-e^{-(\gamma_1+\gamma_2)\zeta}\right].\end{aligned} \tag{S43}$$

Assuming the arteries are positioned symmetrically with respect to the vein (the configuration angles lie on the line of symmetry $\theta_2 = 2\pi - \theta_1$), then $N_{13} = N_{23} = N/2$. Then number of parameters can be reduced to

$$q = 1 - q = 1/2, \quad \mu_1 = \mu_2 = N\mathrm{Da}, \quad \gamma_1 = \gamma_2 = N + 2N_{12} \tag{S44}$$

and hence

$$C_1 = C_2 = g_a(\zeta) = 1 - N\mathrm{Da}\,N_p g_a(1)\zeta, \qquad\qquad C_3 = g_a(\zeta) - N_p g_a(1). \tag{S45}$$

The concentration of the vein can then be written as

$$C_3 = \frac{1 - N_p + N\mathrm{Da}N_p(1-\zeta)}{1 + N\mathrm{Da}N_p}. \tag{S46}$$

For a three vessel cord (S16) becomes

$$\mathcal{E} = \frac{|1 - C_3(0)|}{N_p}, \tag{S47}$$

and by substituting (S46) we obtain the following equation for cord efficiency

$$\mathcal{E} = \frac{1}{|1 + NN_p\mathrm{Da}|}. \tag{S48}$$

A two-vessel cord has the same solution.

### S3. Placental exchange model

As a simple model of placental exchange, we assume that the placental solute, or heat, flux per unit concentration difference, $N_p^*$, can be expressed in terms of the feto-placental Damköhler number $\mathrm{Da}_f$ [62] as

$$N_p^* \approx \mathcal{N}\,B^{-1}\mathcal{L}^* D^* F(\mathrm{Da}_f), \quad \mathrm{Da}_f = \frac{D^*\mathcal{L}^*}{B\,q_f^*} \tag{S49}$$

for some function $F$. Here $\mathcal{L}^*$ is a diffusive length scale specific to terminal villi [7], $\mathcal{N}$ is the number of terminal villi, $B$ is the haemoglobin-facilitated advective factor, $D^*$ is the solute or heat diffusivity in tissue, and $q_f^* \approx Q^*/\mathcal{N}$ is the typical volumetric flow rate per terminal villus. The function $F(\mathrm{Da}_f)$ captures the balance between diffusion-limited and flow-limited transport across the fetal side of the placenta: the maximum placental flux is attained for $F(0) = 1$, when fetal transport is diffusion-limited; in the fetal-flow-limited state, for $\mathrm{Da}_f \gg 1$, the function satisfies $F(\mathrm{Da}_f) \approx \mathrm{Da}_f^{-1}$.

For oxygen exchange in the human placenta at term, we estimate $\mathrm{Da}_f \approx \mathcal{O}\left(10^{-2}\right) \ll 1$, while for heat exchange, $\mathrm{Da}_f \approx \mathcal{O}\left(10^{2}\right) \gg 1$. Our estimates are based on the number of terminal villi $\mathcal{N} \approx \mathcal{O}(10^8)$, approximated as the ratio of the volume of all terminal villi ($\mathcal{O}(10^2\,\mathrm{cm}^3)$, 40% of the total villous tissue volume [9]) to that of a single villous exchange unit ($\mathcal{O}(10^{-6}\,\mathrm{cm}^3)$ [63]), $\mathcal{L}^* \approx \mathcal{O}(10^{-3}\,\mathrm{cm})$ [7], and other parameter values from Table S2.

The umbilical flow-specific placental exchange flux $N_p = N_p^*/Q^*$, predicted by (S49), then reduces to $N_p \sim (\mathcal{N} q_f^*/Q^*)\,\mathrm{Da}_f \approx \mathcal{O}(10^{-2})$ for oxygen and $N_p \sim \mathcal{N} q_f^*/Q^* \approx \mathcal{O}(1)$ for heat. We quantify the uncertainty in the parameter estimates by using an order-of-magnitude variation in $N_p$ to calculate the overall cord transfer efficiency.

### S4. Umbilical blood flow estimation

In order to estimate the cord efficiency across different mammals, measurements of the umbilical blood flow are needed. However, the only available data from the literature were flow measurements from human [24], sheep [25], pony [26] and guinea pig [27]. Few studies have applied allometric scaling to relate body weight and cardiovascular variables across mammals and most of these relationships are described by power-law equations. For example, cardiac output ($\mathrm{cm}^3$/s) was shown to be proportional to body weight (kg) to the 0.74 power [28]; a similar scaling was found between blood flow rate $Q_{MA}$ ($\mathrm{cm}^3$/s) from three major arteries (femoral artery, aorta and common carotid artery) and body weight, $Q_{MA} \propto \mathrm{W}^{0.7-0.8}$ [29]. By fitting a power law to the four measurements available we obtained the following relationship for umbilical blood flow and birth weight $Q^*\,[\mathrm{cm}^3/\mathrm{s}] \approx 1.71\,(\mathrm{BW}\,[\mathrm{kg}])^{\alpha}$, for $\alpha \approx 1.0 \pm 0.14$, which is comparable to the scaling in the major arteries.

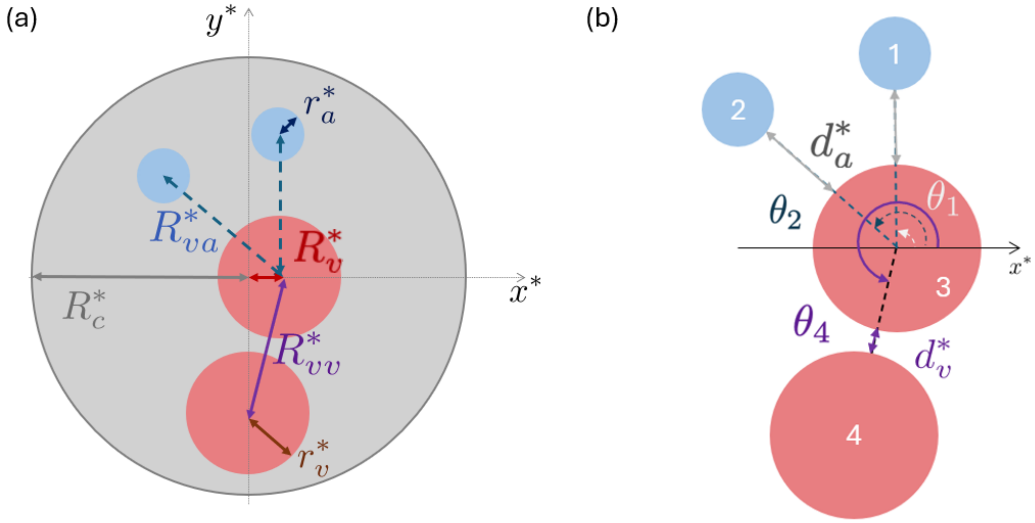


**Figure S1**: **Cross-sectional configuration of the umbilical cord.** (a) A cord's cross-section at $z^* = 0$, with outer radius $R_c^*$. Two veins (red) and two arteries (blue) have radii $r_v^*$ and $r_a^*$ respectively. The centre-to-centre distance between the reference vein and each artery is $R_{va}^*$, and the distance between the centre of the reference vein and the other vein is $R_{vv}^*$. The helical radius $R_v^*$ of the reference vein is the distance between the cord centre and the vein centre. (b) Cross-section in the plane $z^* = 0$ showing vessel positions. The angular position of vessel $i$ is $\theta_i$, measured from the $x^*-$axis. Surface-to-surface separation distances are $d_a^*$ (between the reference vein and each artery) and $d_v^*$ (between veins).

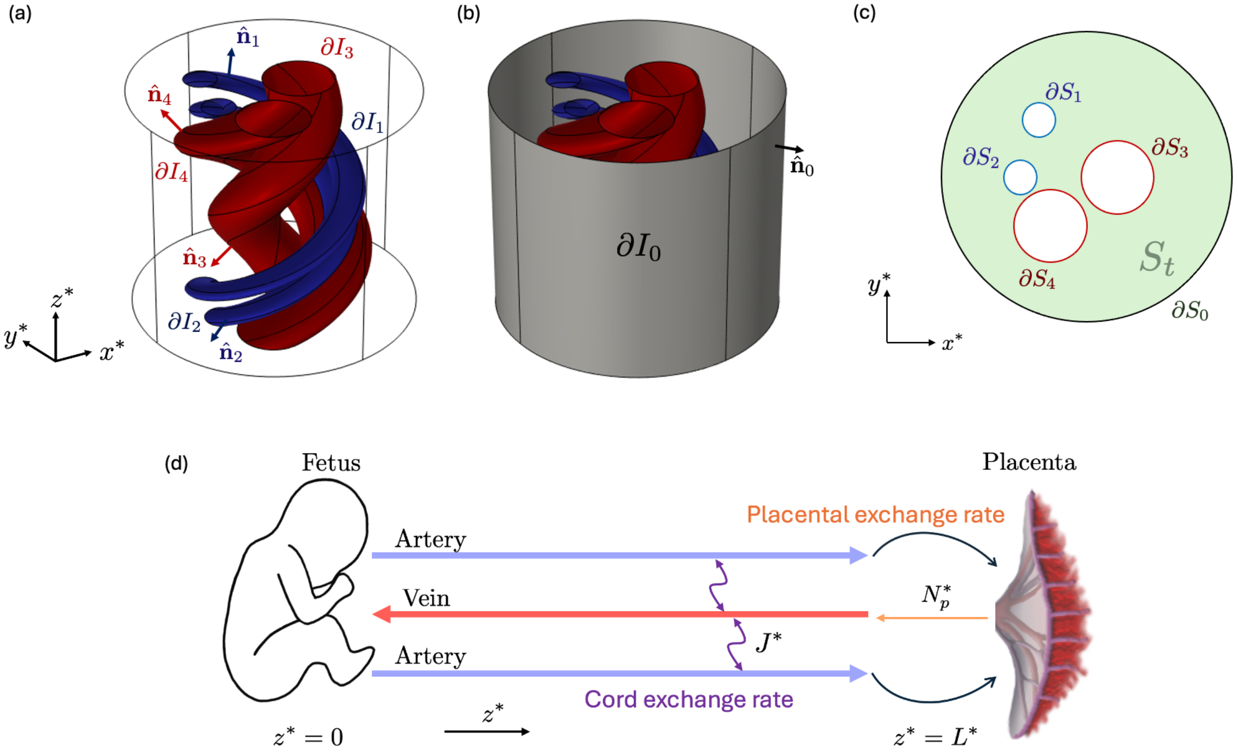


**Figure S2**: **Model geometry and notation**. A four-vessel cord showing (a) $\hat{\mathbf{n}}_i$, the unit normals out of vessel boundaries $\partial I_i$ for $i = 1, 2, 3, 4$; (b) $\hat{\mathbf{n}}_0$, the unit normal out of the cord boundary $\partial I_0 \setminus (I_f \cup I_p)$, (c) the cross-sectional plane of the cord where $\partial S_i$ for $i = 1, 2, 3, 4$ represents the vessel boundary and $S_t$ is the cord tissue. (d) Diagram of the counter-current exchange in a three-vessel cord, illustrating flow directions for each vessel between fetal and placental ends and the corresponding exchange fluxes.

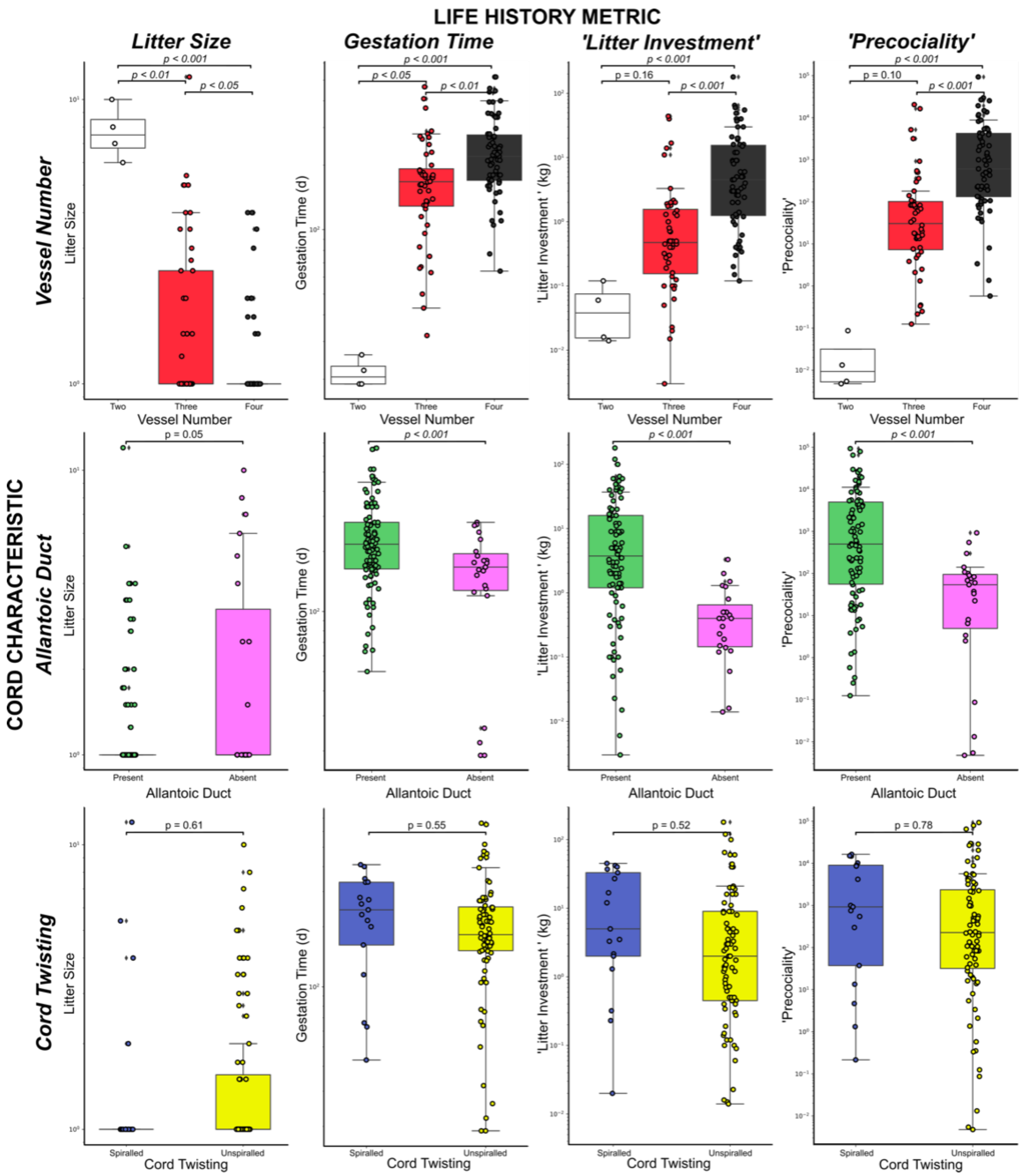


**Figure S3**: **Comparison of umbilical cord characteristics by life history metrics**. Litter investment = (litter size x birth weight). Precociality = ([birth weight x gestation time] / litter size).

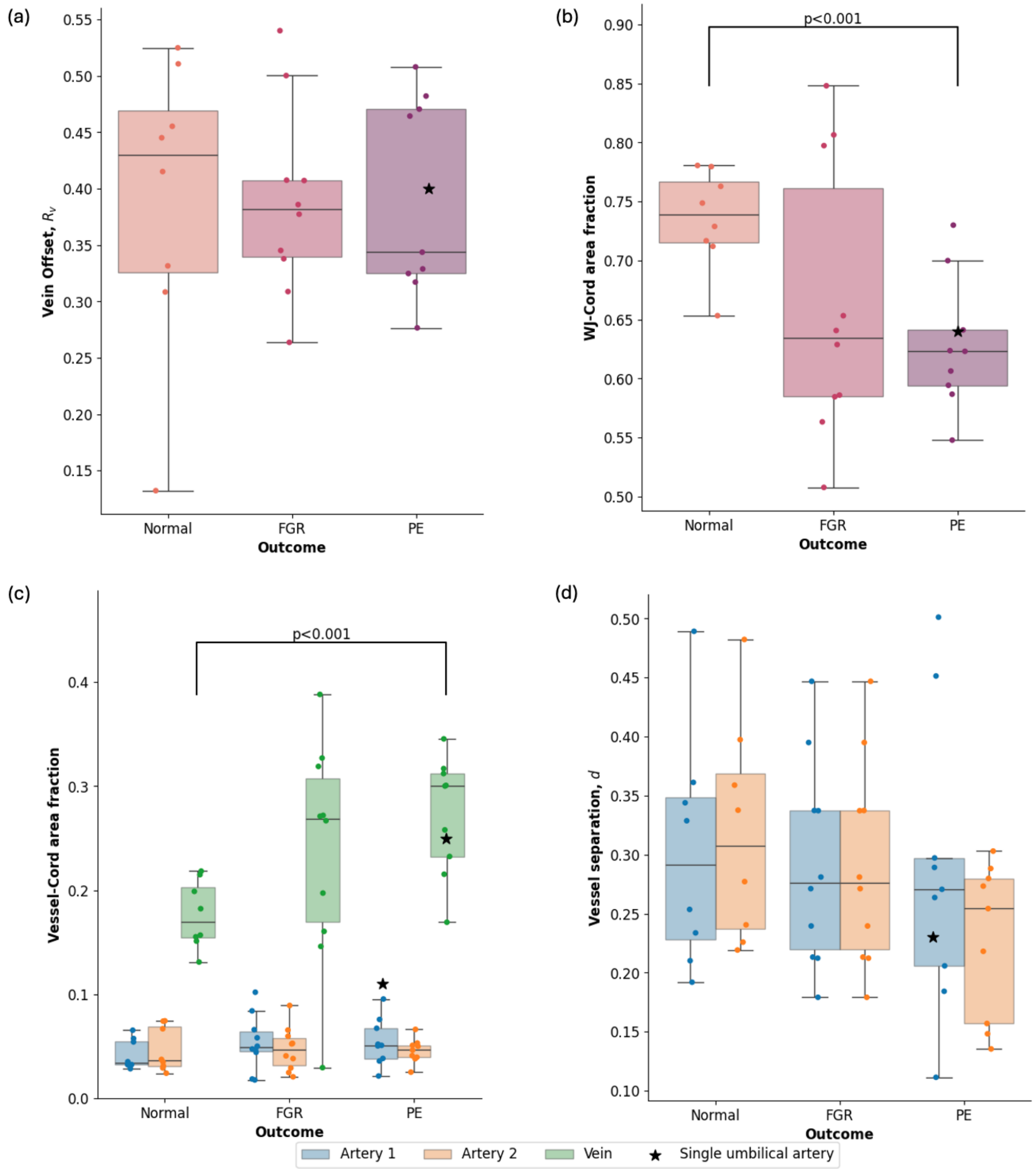


**Figure S4**: **Cross-sectional cord morphometrics**. (a) Vein offset from the cord's centre (scaled with cord radius) for normal and pathological groups; black star marks one case of a single umbilical artery (SUA) in the PE group. (b) Wharton's jelly (WJ) area fraction. (c) Vessel-to-cord area fractions. (d) Artery-vein separation distance (scaled with cord radius). The box displays the median, upper and lower quartiles. The whiskers extend to the farthest point lying within 1.5 times the interquartile range of the sample and points outside this range are considered outliers.

| **Parameter** | **Notation** | **Value** | **Source** |
|---|---|---|---|
| Artery radius | $r_a$ | 0.12 | [64] |
| Vein radius | $r_v$ | 0.25 | [64] |
| Cord radius | $R_c$ | 1 | [64] |
| Vein helical radii | $R_v$ | $0 - 0.17$ | [65] |
| Cord length (human) | $L$ | $42 - 83$ | [9] |
| Pitch (human) | $2\pi/\Omega$ | $1.7 - 12$ | [66] |
| Distance between vessels | $d$ | $0.05 - 0.38$ | Estimated |
| Angle of first artery | $\theta_1$ | $0 - \pi$ | Estimated |
| Angle between arteries | $\Delta\theta \equiv \theta_2 - \theta_1$ | $0.2\,\pi - 1.8\,\pi$ | Estimated |

**Table S1**: **Key dimensionless geometrical parameters of the model**. When the vein helical radius, $R_v$, is zero the vein is straight. The dimensionless values are obtained by scaling the dimensional values from the sources with the cord's radius $R_c^* \simeq 1\,\text{cm}$.

| **Parameter** | **Value** | **Source** |
|---|---|---|
| Oxygen diffusivity, $D^*$ | $2 \times 10^{-5}$ cm$^2$/s | [67] |
| Heat diffusivity, $D^*$ | $1.4 \times 10^{-3}$ cm$^2$/s | [68] |
| Umbilical vein flow rate (human), $Q^*$ | 3 cm$^3$/s | [24] |
| Oxygen advective factor, $B$ | 140 | [58] |
| Heat advective factor, $B$ | 1 | |
| Fetal umbilical oxygen concentration (human), $c_F^*$ | $16 - 28$ mmHg | [69] |
| Maternal placental oxygen concentration (human), $c_M^*$ | 53 mmHg | [7] |
| Feto-maternal temperature difference | $0.5 - 1$°C | [70] |

**Table S2**: **Key physico-chemical parameters of the model**.

| **Quantity** | **2-vessel mouse** | **2-vessel human SUA** | **3-vessel human** | **4-vessel Nile hippo** |
|---|---|---|---|---|
| Representative configuration | | Fig. 4(a) | Fig. 4(b) | Fig. 4(c) |
| Vessel order for $N_{ij}$ | $(a_1, v_1)$ | $(a_1, v_1)$ | $(a_1, a_2, v_1)$ | $(a_1, a_2, v_1, v_2)$ |
| Cord heat-transfer efficiency (S16), $\mathcal{E}_{\text{heat}}$ | 0.15 | 0.82 | 0.75 | 0.95 |
| Cord Damköhler number (S8), $\text{Da}_{\text{heat}}$ | 0.82 | $2.3 \times 10^{-2}$ | $2.3 \times 10^{-2}$ | $2.3 \times 10^{-3}$ |
| Placental exchange parameter (S49), $N_{p,\text{heat}}$ | 3 | 3 | 3 | 3 |
| Cord helicity, $\Omega$ | 0 | 2 | 2 | 2 |
| Mean artery–vein (AV) separation, $d$ | 0.3 | 0.3 | 0.3 | 0.3 |
| Total cross-sectional AV exchange, $N_{\text{tot}}$ | $N_{12} \approx 2.2$ | $N_{12} \approx 3.2$ | $N_{13} + N_{23} \approx 4.9$ | $N_{13}+N_{14}+N_{23}+N_{24} \approx 6.9$ |
| Effective cord coupling, $\text{Da}_{\text{heat}} N_{\text{tot}}$ | 1.8 | $0.74 \times 10^{-1}$ | $1.1 \times 10^{-1}$ | $1.6 \times 10^{-2}$ |
| Pairwise artery–vein flux fractions, $(f_{\alpha\nu})$ | [1] | [1] | $\begin{bmatrix} 1/2 \\ 1/2 \end{bmatrix}$ | $\begin{bmatrix} 0.31 & 0.09 \\ 0.08 & 0.52 \end{bmatrix}$ |
| Exchange fraction: artery 1, $s_{a_1}$ | +1 | +1 | +1/2 | +0.40 |
| Exchange fraction: artery 2, $s_{a_2}$ | — | — | +1/2 | +0.60 |
| Exchange fraction: vein 1, $s_{v_1}$ | −1 | −1 | −1 | −0.39 |
| Exchange fraction: vein 2, $s_{v_2}$ | — | — | — | −0.61 |
| Same-type exchange under $C_{a_i}, C_{v_j} = \text{const}$ | — | — | 0 | 0 |

**Table S3**: **Cross-sectional exchange decomposition for Figs 3 and 4.** The coefficients $N_{ij}$ are the canonical fluxes defined in (S26) and used in (S28); vessel ordering is given in the table. For artery $a_\alpha$ and vein $v_\nu$, define $N_{\text{tot}} = \sum_{\alpha,\nu} N_{a_\alpha v_\nu}$ and $f_{\alpha\nu} = N_{a_\alpha v_\nu}/N_{\text{tot}}$. The signed vessel fractions are $s_{a_\alpha} = \sum_\nu f_{\alpha\nu}$ for heat lost from artery $a_\alpha$ and $s_{v_\nu} = -\sum_\alpha f_{\alpha\nu}$ for heat gained by vein $v_\nu$. In the symmetric three-vessel case, $N_{13} = N_{23} = N_{\text{tot}}/2 \equiv N/2$, while in the four-vessel case $N \equiv N_{\text{tot}} = N_{13} + N_{14} + N_{23} + N_{24}$. Same-type couplings vanish from this decomposition when same-type vessel concentrations are equal; otherwise (S28) must be solved for the axial, concentration-weighted pairwise exchanges.
